# Decoding locomotion from population neural activity in moving *C. elegans*

**DOI:** 10.1101/445643

**Authors:** Kelsey M. Hallinen, Ross Dempsey, Monika Scholz, Xinwei Yu, Ashley Linder, Francesco Randi, Anuj Sharma, Joshua W. Shaevitz, Andrew M. Leifer

## Abstract

The activity of an animal’s brain contains information about that animal’s actions and movements. We investigated the neural representation of locomotion in the nematode *C. elegans* by recording population calcium activity during unrestrained movement. We report that a neural population more accurately decodes locomotion than any single neuron. Relevant signals are distributed across neurons with diverse tunings to locomotion. Two distinct subpopulations are informative for decoding velocity and body curvature, and different neurons’ activities contribute features relevant for different instances of behavioral motifs. We labeled neurons AVAL and AVAR and found their activity was highly correlated with one another. They exhibited expected transients during backward locomotion, although they were not always the most informative neurons for decoding velocity. Finally, we compared population neural activity during movement and immobilization. Immobilization alters the correlation structure of neural activity and its dynamics. Some neurons previously correlated with AVA become anti-correlated and vice versa.

The activity of an animal’s brain contains information about that animal’s actions and movements. We investigated the neural representation of locomotion in the nematode *C. elegans* by recording brain-wide neural dynamics in freely moving animals. We report that a population of neurons more accurately decodes the animal’s locomotion than any single neuron. Neural signals are distributed across neurons in the population with a diversity of tuning to locomotion. Two distinct subpopulations are most informative for decoding velocity and body curvature, and different neurons’ activities contribute features relevant for different instances of behavioral motifs within these subpopulations. We additionally labeled the AVA neurons within our population recordings. AVAL and AVAR exhibit activity that is highly correlated with one another, and they exhibit the expected responses to locomotion, although we find that AVA is not always the most informative neuron for decoding velocity. Finally, we compared brain-wide neural activity during movement and immobilization and observe that immobilization alters the correlation structure of neural activity and its dynamics. Some neurons that were previously correlated with AVA become anti-correlated and vice versa during immobilization. We conclude that neural population codes are important for understanding neural dynamics of behavior in moving animals.

## Introduction

Patterns of activity in an animal’s brain should contain information about that animal’s actions and movements. Systems neuroscience has long sought to understand how the brain represents behavior. Many of these investigations have necessarily focused on single-unit recordings of individual neurons. Such efforts have successfully revealed place cells (***O’Keefe and Dostrovsky, 1971***) and head direction cells (***Taube et al., 1990***; ***Hafting et al., 2005***), for example. But there has also been a long history of seeking to understand how neural populations represent motion (***Georgopoulos et al., 1986***; ***Churchland et al., 2012***; ***Chen et al., 2018***). For example, population recordings from the central complex in *Drosophila* reveal that the animal’s heading is represented in the population by a bump of neural activity in a ring attractor network (***Kim et al., 2017***; ***Green et al., 2017***). As population and whole-brain recording methods become accessible, it has become clear that locomotory signals are more prevalent and pervasive throughout the brain than previously appreciated. For example, neural signals that correlate with rodent facial expression and body motion were recently reported in sensory areas such as visual cortex (***Stringer et al., 2019***) and in executive decision making areas of dorsal cortex (***Musall et al., 2019***).

In *C. elegans*, the field is at a similar inflection point. Locomotion has historically been studied in the worm one neuron at a time using combinations of mutations, ablations (***Gray et al., 2005***), and single neuron recordings of calcium activity (***Arous et al., 2010***; ***Kawano et al., 2011***; ***Piggott et al., 2011***; ***Gordus et al., 2015***; ***Wang et al., 2020a***). Recent work, however, suggests a more important role for neural coding at the level of the population. Population recordings from immobilized animals reveal stereotyped cyclic activity patterns thought to represent global motor commands that account for the majority of the variance in neural dynamics (***Kato et al., 2015***).

In this work we investigate neural representations of locomotion at the population level by recording whole-brain neural activity as the animal crawls freely. We further construct a decoder to predict the animal’s current locomotion from a linear combination of neural activity alone. The performance of the decoder gives us confidence in our ability to find locomotory signals, and allows us to study how those signals are distributed and represented in the brain.

We show that distinct subpopulations of neurons encode velocity and body curvature, and that these populations include neurons with varied tuning. We also find that the decoder relies on different neurons to contribute crucial information at different times. Finally we compared brain-wide neural activity during movement and immobilization and observe that immobilization alters the correlation structure of neural dynamics.

## Results

To investigate locomotory signals in the brain, we simultaneously recorded calcium activity from the majority of neurons in the head of *C. elegans* as the animal crawled freely, ***Figure 1***a-c, (***Nguyen et al., 2016***). The animal expressed the calcium indicator GCaMP6s and a fluorescent protein RFP in the nuclei of all neurons (strain AML310). We report calcium activity as a motion-corrected fluorescence intensity *F*_mc_, described in methods. We measured two features of locomotion: velocity and body curvature related to turning. Velocity is the rate of change of the phase of the body bends propagated along the worm’s body from anterior to posterior derived from an eigenvalue decomposition of the animal’s pose (***Stephens et al., 2008***) and reported in radians per second. We report body bending velocity instead of center of mass velocity, ***Figure 1 - Figure Supplement 1***, because we reasoned that the rate of body bends might more directly reflect the output of the nervous system, as opposed to the center of mass velocity which further depends on mechanical interactions with the substrate. Here we report body curvature as a dimensionless quantity that captures bending in the dorsoventral plane, calculated by projecting the animal’s body posture onto the third principal component of the eigenvalue decomposition. Body curvature captures turning of the animal, but not the small bends required for forward locomotion.

**Figure 1.**
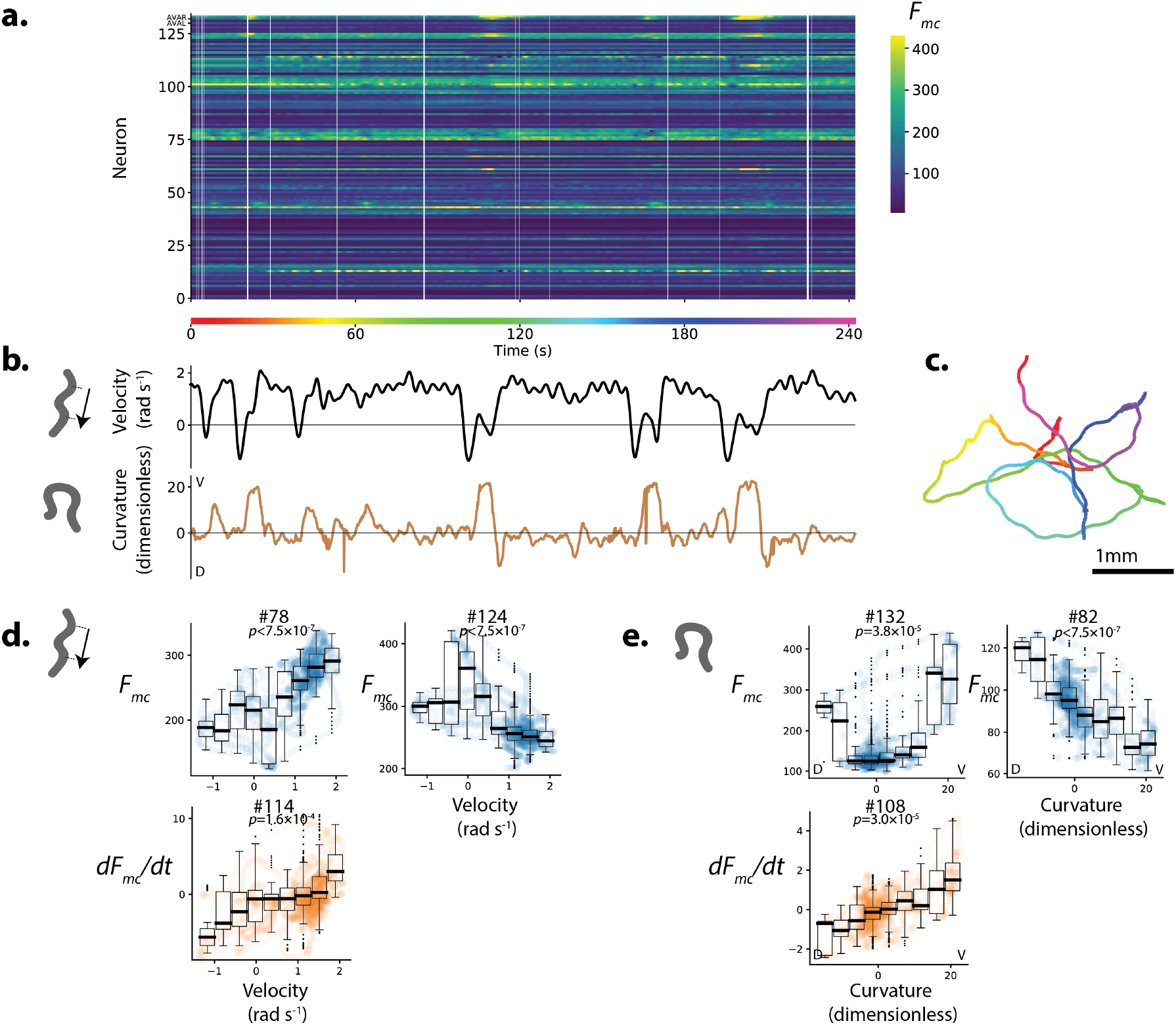
Population calcium activity and tuning of select neurons during spontaneous unrestrained animal movement. Recording AML310_A . a.) Calcium activity of 134 neurons are simultaneously recorded during locomotion. Activity is displayed as motion-corrected fluorescent intensity *F*_*mc*_ . Neurons are numbered according to agglomerative hierarchical clustering. White space indicates time-points where neural tracking failed. b) Body bend velocity and body curvature derived from an eigenvalue decomposition, and c) position on the plate during recording are shown. d.) Example neurons significantly tuned to velocity. Examples are those with the highest Pearson’s correlation coefficient in each category: activity (or its derivative) with positive (or negative) correlation to velocity. P-values are derived from a shuffling procedure that preserves correlation structure. All tuning curves shown are significant at 0.05% after Bonferroni correction for multiple hypothesis testing (*p <* 1.9 × 10^−4^). Boxplot shows median and interquartile range. e) Example neurons highly tuned to curvature were selected similarly. No neurons with negative *dF* /*dt* tuning passed our significance threshold. **Figure 1–Figure supplement 1**. Comparison of center-of-mass velocity and eigenvalue decomposition-derived velocity.

### A diversity of neural tuning to behavior exists in the population

We found multiple neurons with calcium activity significantly tuned to either velocity or curvature (***Figure 1***). Some neurons were more active during forward locomotion while others were more active during backward locomotion (***Figure 1***d). Similarly some neurons were active during dorsal bends and others during ventral bends (***Figure 1***e). In some cases, the derivative of the activity was also significantly correlated with features of locomotion. Significance was calculated using a shuffe and a Bonferroni correction was applied. The existence of such neural signals correlated with these behaviors is broadly consistent with single-unit or sparse recordings during forward and backward locomotion (***Arous et al., 2010***; ***Kawano et al., 2011***; ***Gordus et al., 2015***; ***Shipley et al., 2014***; ***Kato et al., 2015***; ***Wang et al., 2020b***) and turning (***Kocabas et al., 2012***; ***Donnelly et al., 2013***; ***Wang et al., 2020b***).

AVA’s activity during population recordings is consistent with prior reports We labeled the neurons AVAL and AVAR using blue fluorescent protein (BFP) which is spectrally separated from the other two colors we use for neuron localization and activity (strain AML310) to unambiguously identify this well-characterized neuron pair during population calcium imaging recordings, see ***Figure 2***a. These two neurons, called AVA, are a bilaterally symmetric pair with gap junctions between them that have been shown to exhibit large calcium transients that begin with the onset of backward locomotion, peak around the end of backward locomotion during the onset of forward locomotion, and then slowly decay (***Arous et al., 2010***; ***Kawano et al., 2011***; ***Shipley et al., 2014***; ***Gordus et al., 2015***; ***Kato et al., 2015***). Our measure of AVA’s activity, recorded simultaneously with 131 other neurons during unrestrained movement, is consistent with prior recordings where AVA was recorded alone. We note that single-unit recordings of AVA used in previous studies lacked the optical sectioning needed to resolve these neurons separately. Here we resolve both AVAL and AVAR and find that their activities are similar to one another, and they both exhibit the expected transients timed to backward locomotion, ***Figure 2***b. Signal-to-noise in AVAR is higher than AVAL because in this recording AVAR lies closer to the imaging objective, while AVAL is on the opposite side of the head and therefore must be imaged through the rest of the brain. The similarity we observe between activities of AVAL and AVAR demonstrates our ability to simultaneously record neural activity accurately from across the entire brain.

**Figure 2.**
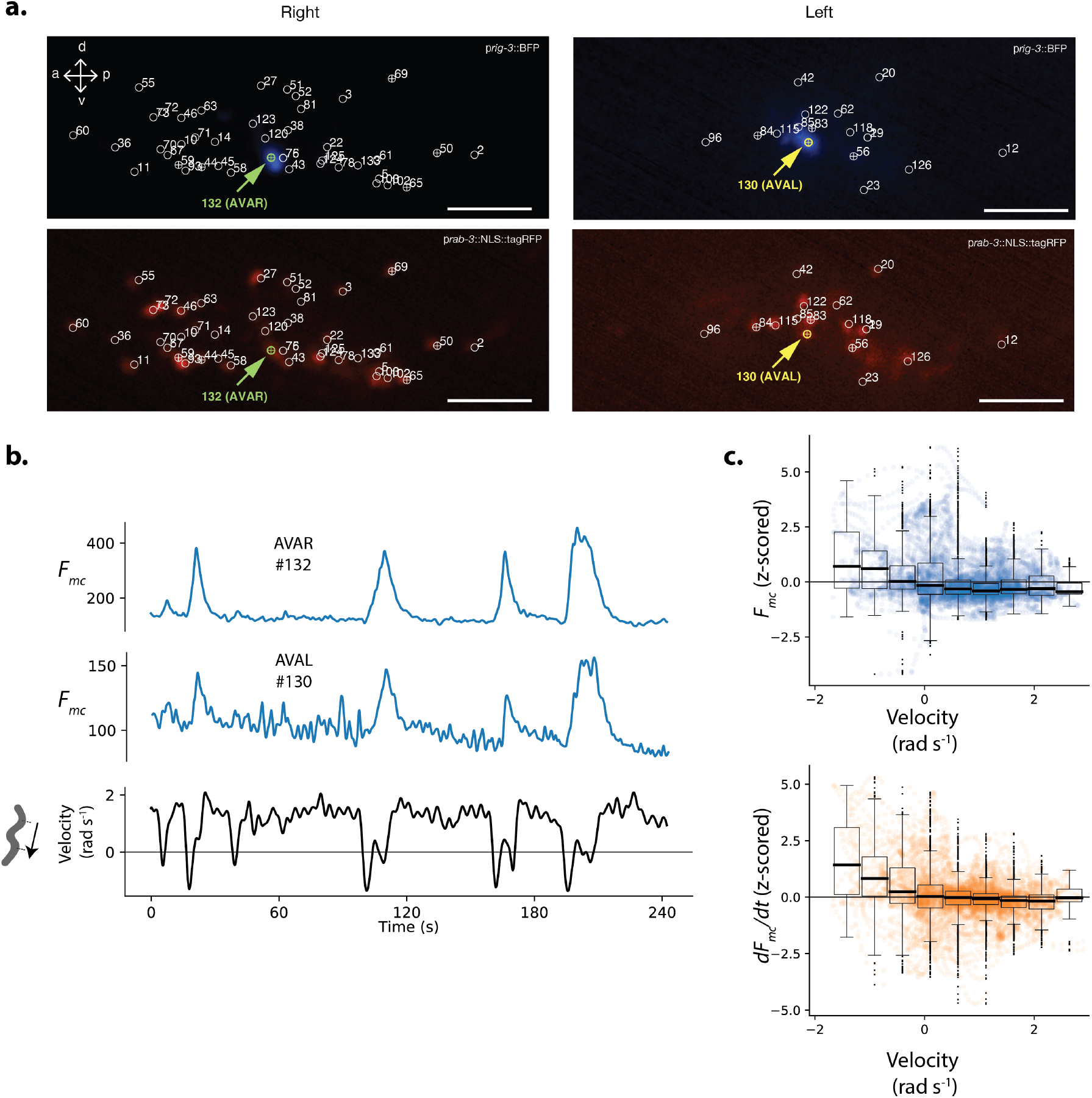
Neuron pair AVA is active during backward locomotion and exhibits expected tuning during freely moving population recordings. a.) AVAR and AVAL are labeled by BFP under a *rig-3* promoter in strain AML310. Two optical planes are shown from a single volume recorded during free movement. Planes are near the top and bottom of the optical stack, corresponding to the animals’ extreme right and left. The recording is the same as in ***Figure 1***. Top row shows BFP. Bottom row shows RFP in the nuclei of all neurons. Segmented neurons centered in the optical plane are labeled with *EB*, while neurons from nearby optical planes are labeled with o. Arrow indicates AVAR or AVAL. Numbering corresponds to ***Figure 1***a. b.) Calcium activity of AVAR and AVAL during locomotion in recording AML310_A, same as in ***Figure 1***. c.) Aggregate tuning of AVA across four individuals (7 neurons). Boxplot shows median and interquartile range.

We recorded from three additional animals and identified AVA neurons in each. The temporal derivative of AVA’s activity has previously been shown to correlate with velocity over the range of negative (but not positive) velocities (***Kato et al., 2015***). Consistent with these reports, the derivative of AVA’s activity, *dF*_mc_/*dt*, aggregated across the four population recordings has a negative correlation to velocity over the range of negative velocities, ***Figure 2***c.

In our exemplar recording, AVA’s activity (not its temporal derivative) also correlates with body curvature (***Figure 1***e, neuron #132). Correlation to curvature likely arises because our exemplar recording includes many long reversals culminating in deep ventral bends called “omega turns,” that coincide in time with AVA’s peak activity. Taken together, AVA’s activity simultaneously recorded from the population is in agreement with prior reports where AVA activity was recorded alone.

### Population decoder outperforms best single neuron

AVA’s activity is related to the the animal’s velocity, but its activity alone is insuffcient to robustly decode velocity. For example, AVA is informative during backward locomotion, but contains little information about velocity during forward locomotion, ***Figure 2***c. To gain reliable information about velocity, the nervous system will need more than just the activity of AVA. In primate motor cortex, for example, linear combinations of activity from the neural population provides more information about the direction of a monkey’s arm motion during a reach task than a single neuron (***Georgopoulos et al., 1986***). We wondered whether activity of the neural population might be more informative of the worm’s locomotion than an individual neuron.

We constructed a population decoder using linear regression with regularization to find neural weights that relate recorded population activity to each behavior: velocity and curvature. Ridge regression (***Hoerl and Kennard, 1970***) was performed on 60% of the recording (training set) and the decoder was evaluated on the remaining 40% (held-out test set, shaded green in ***Figure 3***a,c). Cross-validation was used to set hyper-parameters (described in methods). Our decoder assigned two regression coeffcients to each neuron, one weight for activity and one for its temporal derivative. We compared performance of the population decoder to that of the most correlated single neuron or its derivative. Performance is reported as a coeffcient of determination on the mean-subtracted held out test set 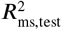.For the exemplar recording shown in ***Figure 1*** and ***Figure 2***a-b the population performed better on the held-out-test set than the most correlated single neuron (or its temporal derivative) for both velocity and body curvature, see ***Figure 3***. For velocity, population performance was 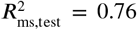 compared to 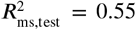 for the best single neuron; and for curvature population performance was 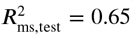 compared to 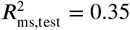 for the best sin-gle neuron. Red arrows in ***Figure 3*** highlight striking behavior features that the best single neuron misses but that the population decoder captures. We also explored alternative population models, including both linear and non-linear models with different features and cost penalties, ***Figure 3 - Figure Supplement 3***. Of the populations models we tried, the model used here was one of the simplest and also had the best mean performance at decoding velocity across all recordings.

**Figure 3.**
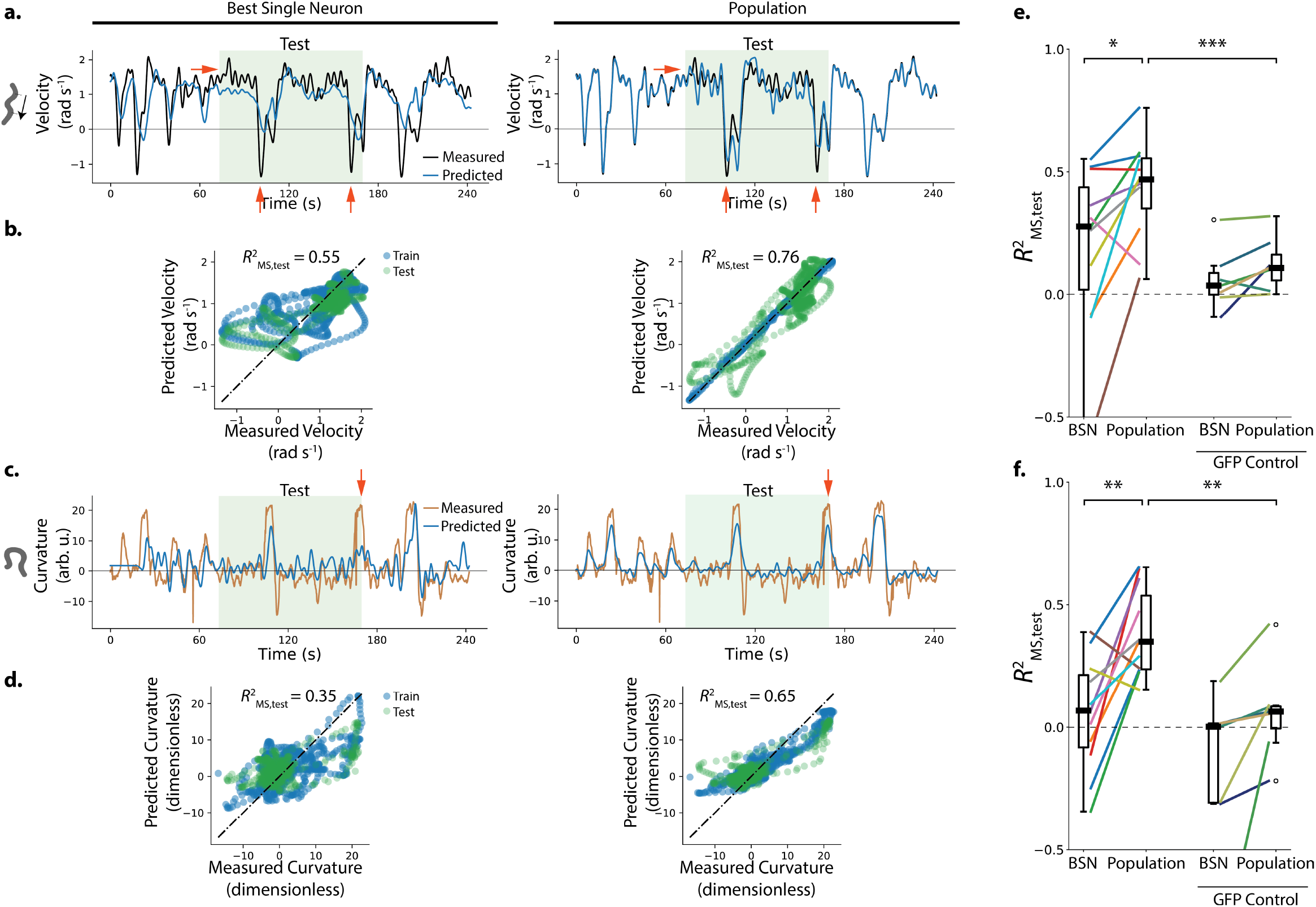
Population neural activity decodes locomotion. a-d.) Performance of the best single neuron (BSN) is compared to a linear population model in decoding velocity and body curvature for the exemplar recording AML310_A shown in ***Figure 1***. a.) Predictions on held-out test set are compared to measured velocity. Light green shaded region indicates held-out test set. Red arrows indicate examples of features that the population captures better than the BSN. b.) Performance is reported as a coeffcient of determination 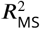 evaluated on the mean-subtracted held-out test data (green points). c,d) Model predictions are compared to measured curvature. e) Performance of velocity decoding is shown for recordings of *n* = 11 individuals (strain AML310 and AML32) and for recordings of *n* = 7 GFP control animals lacking a calcium indicator (strain AML18). Two-sided Wilcoxon rank test is used to test significance of population performance compared to BSN, *p* = 0.014. Welch’s unequal variance t-test is used to test significance of population performance compared to GFP control, *p* = 8.0 × 10^−4^ f) Performance of curvature decoding is shown for all recordings. Each recording is colored the same as in e. *p* = 4.8 × 10^−3^ and *p* = 3.8 × 10^−3^ for comparisons of population performance to that of BSN, and GFP control, respectively. **Figure 3–Figure supplement 1**. Performance correlates with maximal GCaMP Fano Factor, a metric of signal. **Figure 3–Figure supplement 2**. Neural activity and behavior for all recordings. **Figure 3–Figure supplement 3**. Alternative population models.

Activity was recorded from a total of 11 animals during unrestrained motion and the linear population model was used to decode each recording (n=7 recordings of strain AML32; n=4 recordings of strain AML310, also shown in ***Figure 2***c). The population significantly outperformed the best single neuron at decoding the held-out portions of the recordings for both velocity and curvature (*p <* 0.05 two-sided Wilcoxon rank test).

There was large worm-to-worm variability in the performance of the decoders. Performance across recordings correlated with one metric of the signal in our recordings, the maximal Fano factor across neurons of the raw time-varying GCaMP fluorescence intensity,

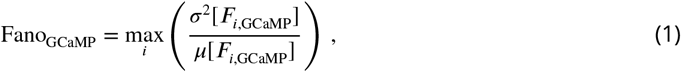

where max_*i*_ indicates the maximum over all neurons in the recording, and *σ*,^2^ and *µ* are the variance and mean respectively of the raw GCaMP activity of the neuron, see ***Figure 3 - Figure Supplement 1***. The recording with the highest Fano_GCaMP_ performed best at decoding velocity and curvature. This suggests that variability in performance may be due in part to variability in the amount of neural signal in our recordings.

In some recordings where the population outperforms the best single neuron, it does so in part because the population decodes a fuller range of the animal’s behavior compared to the best single neuron. Recording AML32_A shows a striking example: the best single neuron only captures velocity dynamics for negative velocities. The population decoder, by contrast, captures velocity dynamics during both forward and backward locomotion during the held-out test set, and covers a larger fraction of the held-out velocity range, see ***Figure 4***.

**Figure 4.**
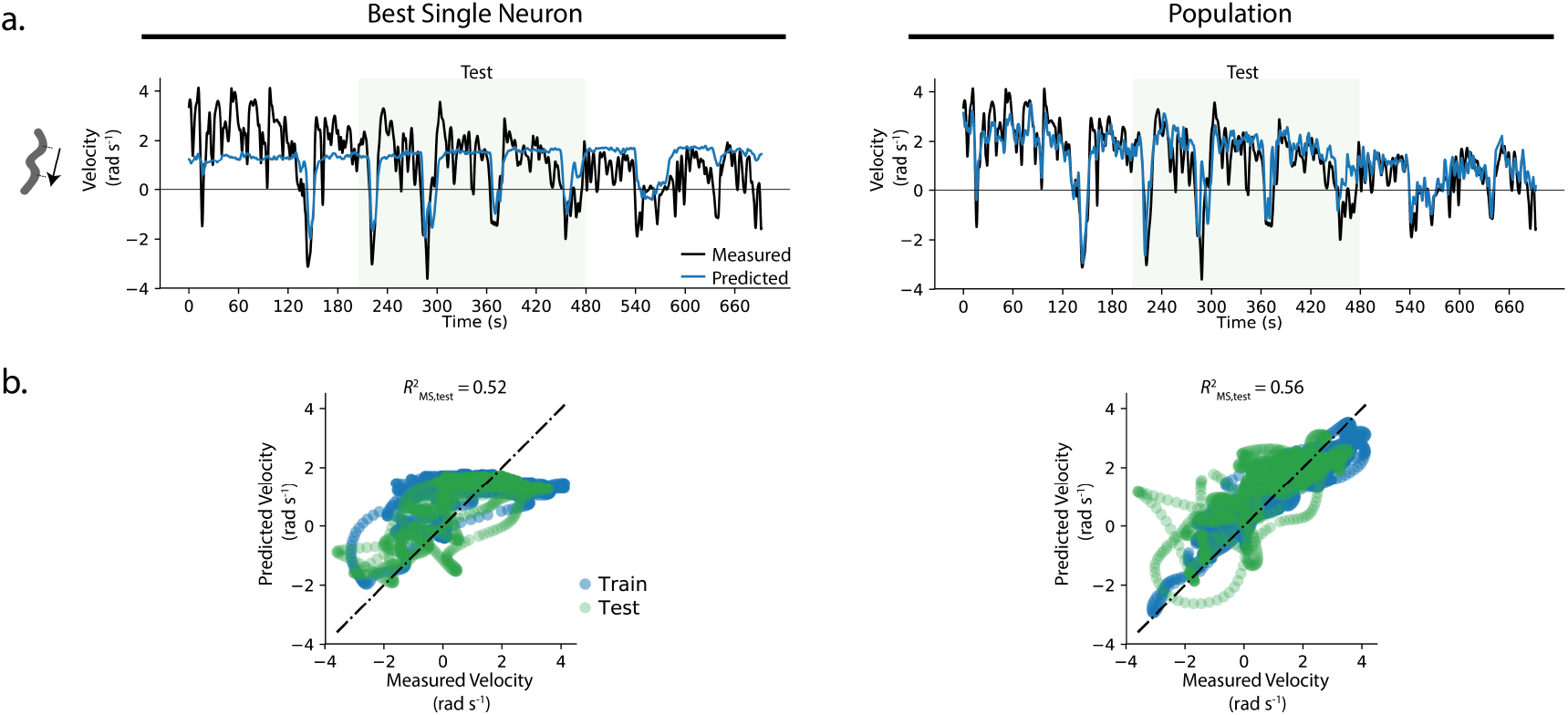
Example where population decoded a fuller range of animal behavior. a.) The decoding from the best single neuron and the population model are compared to the measured velocity for example recording AML32_A . b.) Predictions from the best single neuron saturate at a velocity of 2 rad s^−1^.

Motion artifact is of potential concern because it may resemble neural signals correlated to behavior (***Nguyen et al., 2016***; ***Chen et al., 2013***). For example, if a neuron is compressed during a head bend, it may increase local fluorophore density causing a calcium-independent increase in fluorescence that would erroneously appear correlated with head bends. We address this concern in all our recordings by extracting a motion corrected calcium signal derived from a comparison of GCaMP and RFP dynamics in the same neuron. All strains in this work express a calcium insensitive RFP in every neuron in addition to GCaMP. Motion artifacts should affect both fluorophores similarly. Therefore, the motion correction algorithm extracts only those GCaMP dynamics that are independent of the RFP timeseries, and it rejects dynamics common to both (details in methods). To validate our motion correction, and to rule out the possibility that our decoder primarily relies on non-neural signals such as those from motion artifact, we recorded from control animals lacking calcium indicators. These animals expressed GFP in place of GCaMP (7 individuals, strain AML18, RFP was also expressed in all neurons). GFP emits a similar range of wavelengths to GCaMP but is insensitive to calcium. Recordings from these control animals were subject to similar motion artifact but contained no neural activity because they lack calcium sensors (***Figure 3 Figure Supplement 2***). Recordings from GFP control animals were subject to the same motion correction as GCaMP animals. For both velocity and curvature, the average population model performance was significantly worse at decoding calcium-insensitive GFP control recordings than the calcium-sensitive GCaMP recordings (***Figure 3e-f***, median performance 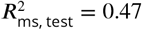 for GCaMP compared to 0.11 for GFP control at decoding velocity, and median performance 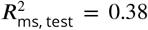 for GcaMP compared to 0.06 for GFP control for curvature, *p <* 0.01 Welch’s unequal variance test), suggesting that the decoder’s performance relies on neural signals.

### Population code for locomotion

The distribution of weights assigned by the decoder provides information about how behavior is represented in the brain. Each neuron was assigned one weight for its activity *W*_*F*_ and another for the temporal derivative of its activity 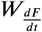. In the exemplar recording from ***Figure 1***, weights were distributed roughly evenly between positive and negative and were well approximated by a single Gaussian distribution centered at zero, see ***Figure 5ab***. The decoder relied on both neural activity and its temporal derivative and assigned neural weights roughly evenly between the two. The weight assigned to a neuron’s activity *W*_*F*_ was not correlated with the weight assigned to its temporal derivative 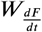 (***Figure 5-Figure Supplement 1***).

**Figure 5.**
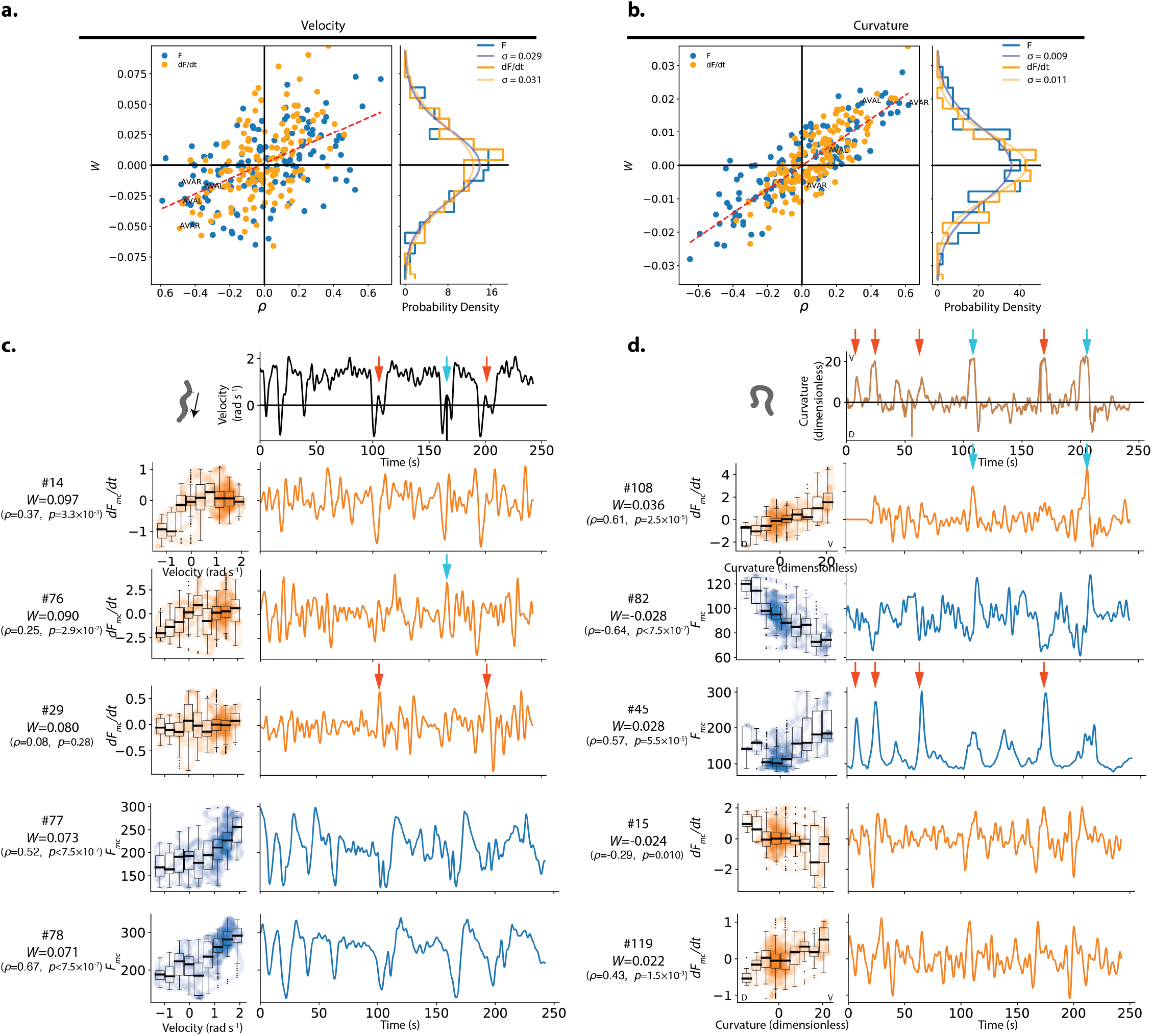
Weights assigned to neurons by the population model in the exemplar recording, and their respective tuning. a.) The weight *W* assigned to each neuron’s activity (*F*_mc_) or its temporal derivative (*dF*_mc_/*dt*) by the velocity population decoder is plotted against its Pearson’s Correlation coefficient *ρ* which characterizes its tuning to velocity. Recording AML310_A is shown, same as in ***Figure 1***. Dashed red line shows line of best fit. Right panel shows the observed distribution of weights. A zero-mean Gaussian with standard deviation set to the empirically observed standard deviation is also shown. b.) Same as in a, but for curvature. c.) Tuning and activity of the top five highest amplitude weighted neurons is shown. Activity of each neuron is time aligned to the observed behavior (top row). Neurons are labeled corresponding to their number in the heatmap in ***Figure 1***. Their weight *W* in the decoder and the Pearson’s correlation coefficient to velocity are also listed. Red and cyan arrows highlight peaks in the temporal derivative of neuron #29’s activity and in neuron #76’s activity that contribute to predicting the slight increase in velocity observed during complementary instances when the animal initiates a deep ventral bend. Y- and X-axes labels and scales are preserved within individual rows and columns, respectively. d) Same as c but for curvature. Red and cyan arrows show two sets of deep ventral bends that are captured by two neurons. **Figure 5–Figure supplement 1**. Comparison of weights assigned to a neuron’s activity versus its temporal derivative. **Figure 5–Figure supplement 2**. Comparison of weights assigned for decoding velocity vs decoding curvature.

In general, a neuron ’s weight is correlated with its Pearson’s correlation coeffcient to behavior (trend visible in ***Figure 5ab***), but individually, many neurons are weighted in non-intuitive ways suggesting that the linear decoder in our model involves more than merely assigning high weights to those neurons with the highest correlation to behavior. For example, the most negatively weighted neuron for velocity had a slightly positive tuning (Pearson’s correlation *ρ >* 0, ***Figure 5***a). Similarly the temporal derivative of neuron #29, discussed in detail below (***Figure 5***b), had the third highest magnitude weight for velocity but shows no obvious tuning (correlation coeffcient to velocity is *ρ* = 0.25, which is below our threshold for statistical significance). That some prominently weighted neurons exhibit little tuning to locomotion, or have opposite signed tuning than that of their assigned weight suggests that the decoder relies in part on aspects of neural activity that are not easily captured by a tuning curve. This prompted us to investigate neural weighting further.

We inspected the activity traces of the top five weighted neurons in our exemplar recording (***Figure 5***c,d). Some highly weighted neurons had activity traces that appeared visually similar to the animal’s locomotory trace for the duration of the recording (e.g. #77 and #78 for velocity). But other highly weighted neurons had activity traces that seemed to only match specific features of the locomotory behavior and only then for specific brief times or for specific instances of a behavior motif. For example, the temporal derivative of the activity of neuron #29 contributes a subtle but distinct peak to velocity at times 105 s and 200 s when the animal slows down its reversal before executing a bend and initiating forward locomotion, but not during the similar behavior at time 160 s (***Figure 5***c). The features contributed by neuron #29 are missing from the other neurons in the top five. Conversely, neuron #76 in the top five contributes a prominant peak of activity that matches the locomotry feature at 160 s that was absent from neuron #29. Similarly for curvature, neuron #45 has large sharp peaks of activity during four of six deep ventral bends, but only modest transients during the remaining two ventral bends at approximately 110s and 210 s (***Figure 5***d). Conversely neuron #108 had pronounced peaks of the temporal derivative of its activity at the deep ventral bends at 110 s and 210 s but not at the other four ventral bends. We note this phenomonenon was present in the test set as well as the training set.

We conclude that there are at least two types of neural signals that the linear model weighs highly for decoding locomotion. One type are neural signals that are consistently highly tuned to locomotion (e.g neuron #78). The other type are neural signals with pronounced activity relevant only for particular instances of behavioral motifs (e.g #45, and #108). For the latter, the decoder appears to weight multiple neurons highly to capture all instances of the behavior.

#### Majority of decoder’s performance is provided by a subset of neurons

We sought to better understand how information used by this particular model is distributed among the population. We wondered how many neurons the model relies upon to achieve most of its performance. The magnitude of a neuron’s weight in the population model is indicative of its relative usefulness in decoding locomotion. Therefore we investigated performance of a restricted population model that had access to only those *N* neurons that were most highly weighted by the full model. We sequentially increased the number of neurons *N* and evaluated the partial model performance (***Figure 6*** - Video Supplement 1). In this way we estimated the number of neurons needed to first achieve a given performance (***Figure 6***a). Because we were interested in probing the particular successful set of weights that the model had found, we constrained the relative weights of neurons in the partial model to match those of the full model. We note that adding a neuron gave the model access to both that neuron’s activity and its temporal derivative. We define the number of neurons needed to first achieve 90% full model performance as the *N*_90_ and use this value as an estimate of the number of important neurons for decoding. For the exemplar recording `AML310_A`, 90% of the model’s performance was achieved when including only 28 neurons for velocity, and only 3 neurons for curvature.

**Figure 6.**
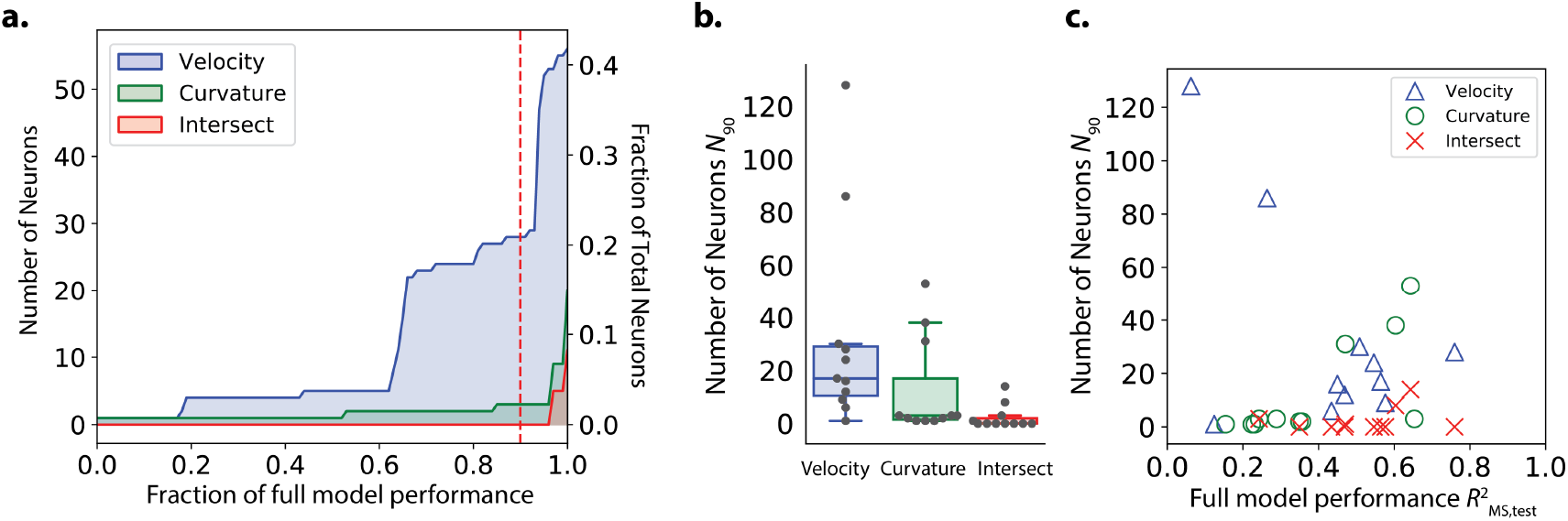
Number of neurons needed by the model to decode velocity and curvature. a.) The minimum number of neurons needed for a restricted model to first achieve a given performance is plotted from recording AML310_A in ***Figure 1***. Performance is reported separately for velocity (blue) and curvature (green) calculated on the held-out test set. Intersect refers to the intersection of the set of neurons included in both partial models (velocity and curvature) for a given performance. Red dashed line, *N*_90_, indicates number of neurons needed to achieve 90% of full model performance. b.) *N*_90_ is computed for velocity and curvature for all recordings. The number of neurons present in both populations at 90% performance level (intersection) is shown. Box shows median and interquartile range. c) *N*_90_ for all recordings is shown plotted verses the performance of the full population velocity or curvature decoder, respectively. Number of intersection neurons (red ‘x’) is plotted at the higher of either the velocity or curvature’s performance. **Figure 6–video 1**. Animation showing partial model performance as neurons are added, corresponding to ***Figure 6***a. Top panel shows performance. Bottom left shows measured velocity (black) and decoded velocity (blue). Gray shading indicates test set. Bottom right shows measured velocity compared to decoded velocity for training (blue) and test set (green).

Across all recordings we saw large variability in the number of important neurons *N*_90_ (***Figure 6***b and ***Table 1***) with a median of 33 neurons for velocity and 13 for curvature. Decoder-recording instances that exhibited high full model performance 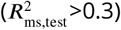 always had less than 55 neurons in each velocity and curvature sub-populations (***Figure 6***c). By comparison, our recordings contained a median total of 121 neurons. On average, the decoder relies on less than a third of the neurons in a recording to achieve the majority of its decoding performance.

**Table 1.** Number of neurons needed to achieve 90% of full model performance, *N*_90_, reported as (median ± standard deviation), across all 11 recordings.

| Velocity $N_{90}$ | Curvature $N_{90}$ | Intersection $N_{90}$ | Total Recorded |
| --- | --- | --- | --- |
| 33 ± 37 | 13 ± 18 | 2 ± 4 | 121 ± 12 |

#### Largely distinct sub-populations contain information for velocity and curvature

We wondered how a neuron’s role in decoding velocity related to its role in decoding curvature. In exemplar recording AML310_A, there was no obvious population-wide trend between the magnitude of a neuron’s weight at decoding velocity and the magnitude of its weight at decoding curvature for either *F*, *dF* /*dt* or both, see ***Figure 5 - Figure Supplement 2***. Furthermore, there was no overlap between the *N*_90_ = 23 neurons needed to achieve 90% of full model performance at decoding velocity and the *N*_90_ = 3 neurons needed for curvature in this recording, see ***Figure 6***a. Across all recordings only 2 ± 4 (median ± std) neurons were included in both *N*_90_ for the velocity and curvature sub-populations, labeled “intersect” neurons in ***Figure 6***bz,c and ***Table 1***. Taken together, this suggests that largely distinct sub-populations of neurons in the brain contain the majority of information important for decoding velocity and curvature.

### Immobilization alters the correlation structure of neural dynamics

Brain-wide activity during immobilization has previously been studied to gain insights into neural dynamics of *C. elegans* locomotion (***Kato et al., 2015***). We therefore investigated the effect of immobilization on neural dynamics. We recorded population activity from a freely moving animal crawling in a microfluidic chip and then immobilized that animal partway through the recording by delivering the paralytic levamisole, as has been used previously (***Gordus et al., 2015***; ***Kato et al., 2015***). Neural dynamics from the same neurons in the same animal were therefore directly compared during movement and immobilization, ***Figure 7***.

**Figure 7.**
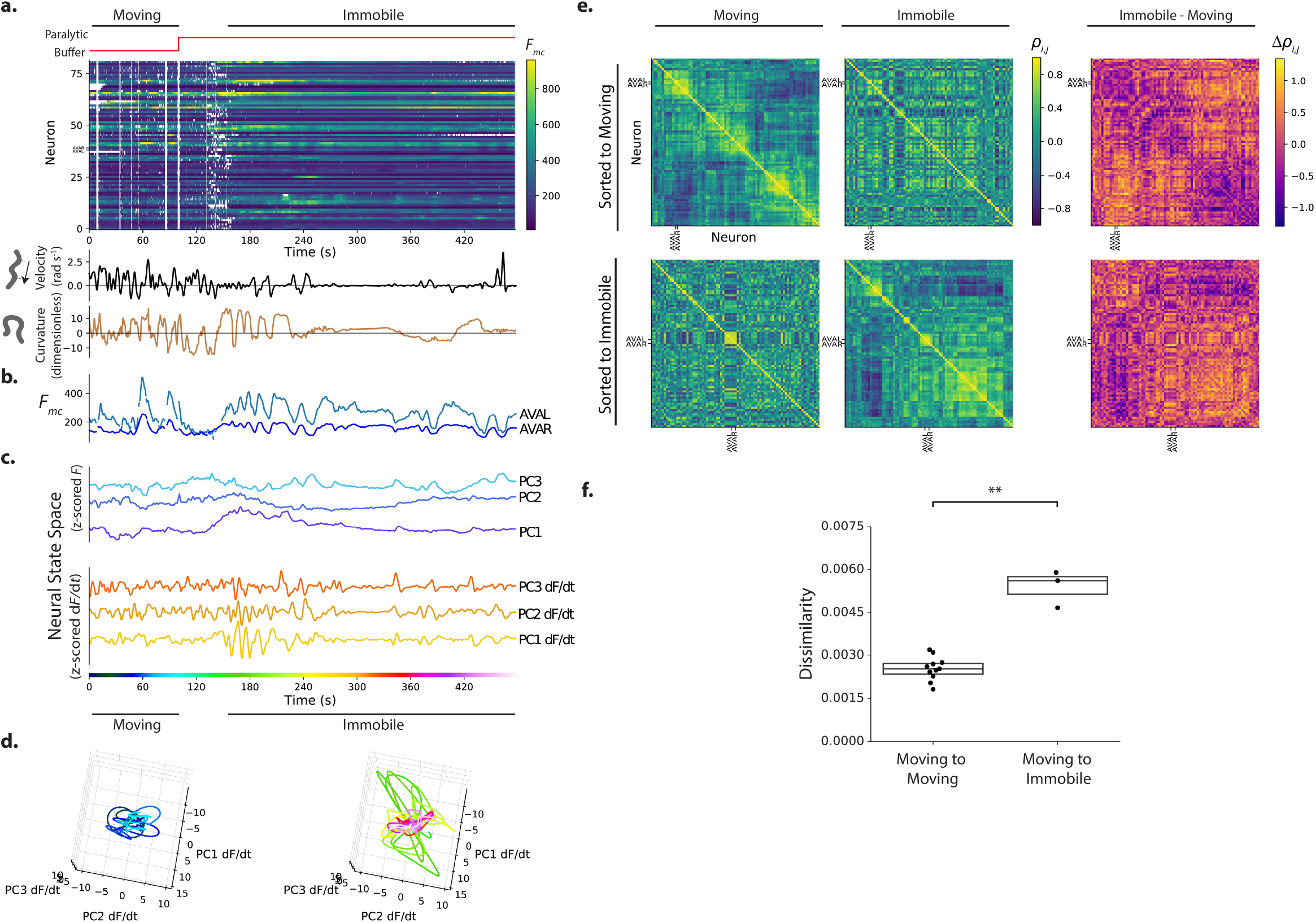
Immobilization alters the correlation structure of neural activity. a). Calcium activity is recorded from an animal as it moves and then is immobilized with a paralytic drug, recording AML310_E . b.) Activity of AVAL and AVAR from (a). c.) Population activity (or its temporal derivative) from (a) is shown projected onto its first three PCs, as determined by only the immobilized portion of the recording d.) Neural state space trajectories from (c) are shown split into moving and immobile portions, color coded by time. Scale and axes are same. e.) Pairwise correlations of neural activity *p*_*i,j*_ are shown as heatmaps for all neurons during movement and immobilization, sorted via clustering algorithm. Top row is sorted to movement, bottom row is sorted to immobilization. f) Dissimilarity between correlation matrices for moving and immobile portions of a recording are shown compared to the dissimilarity observed between correlation matrices taken at similar time windows within moving-only recordings. Dissimilarity is 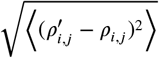. Dissimilarity was measured in 3 moving-immobile recordings with paralytic and 11 moving-only recordings. *p* = 9.9 × 10^−3^, Welch’s unequal variance t-test. Boxes show median and interquartile range. **Figure 7–Figure supplement 1**. Example from additional moving-to-immobile recording. **Figure 7–Figure supplement 2**. Immobile-only recording.

**Figure 8.**
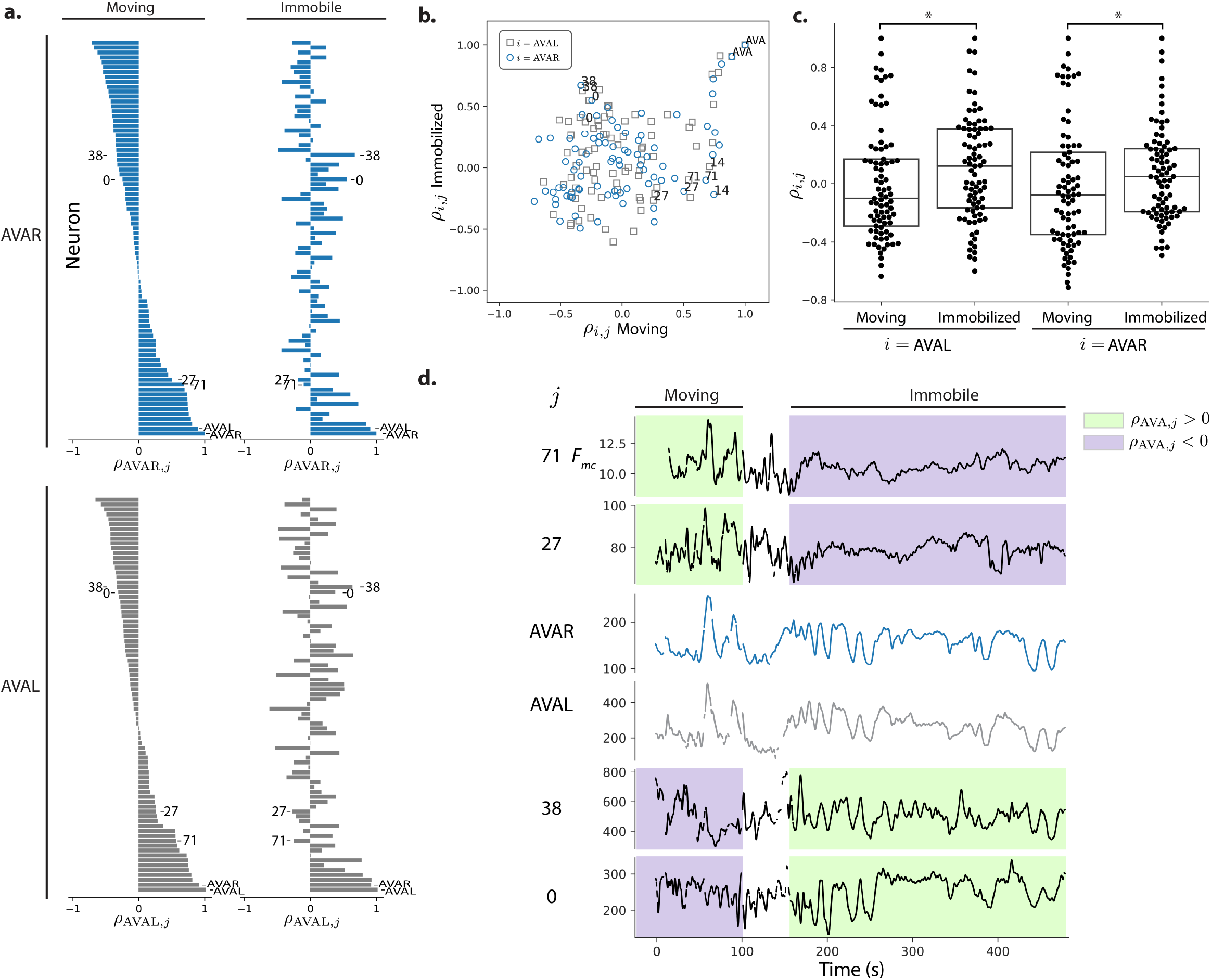
Correlations with respect to AVAL and AVAR during movement and immobilization. a.) The pearson’s correlation of each neuron’s activity to AVAR and AVAL is shown during movement and immobilization. Selected neurons are numbered as in ***Figure 7*** (same recording, AML310_E). Neurons are sorted according to their correlation during movement. b.) Scatter plot shows relation between a neuron’s correlation to AVA during movement and its correlation during immobilization. Gray squares and blue circles indicate correlation to AVAL and AVAR, respectively. c.) On average, neurons become more positively correlated to AVAL and AVAR, *p* = 0.028 and *p* = 0.038, respectively, Welch’s unequal variance t-test. Box shows median and interquartile range. d.) Activity traces of selected neurons are shown time aligned to AVA. Green and purple shading indicate positive or negative correlation to AVA, respectively.

Immobilization changed the correlation structure of neural activity. Clusters of neurons that had been correlated with one another during movement were no longer correlated during immobilization (see ***Figure 7***e, top row, blocks of contiguous yellow on the diagonal during movement that are absent or disrupted during immobilization). Notably, many neurons that had been only weakly correlated or even anti-correlated during movement became correlated with one another during immobilization forming a large block (***Figure 7***e, bottom, large contiguous yellow square that appears on the lower right along the diagonal during immobilization).

To further quantify the change in correlation structure, we defined a dissimilarity metric, the root mean squared change in pairwise correlations 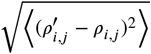, and applied it to the correlation matrices during movement and immobilization within this recording, and also to two additional recordings with paralytic. As a control, we also measured the change in correlation structure across two similar time windows in the 11 freely moving recordings. The change in correlations from movement to immobilization was significantly larger than changes observed in correlations in the moving-only recordings (*p* = 9.9 × 10^−3^, Welch’s unequal variance t-test) see ***Figure 7***f and methods. This suggests that immobilization alters the correlation structure more than would occur by chance in a freely moving worm.

We next inspected the neural dynamics themselves (***Figure 7***a, c). Low-dimensional stereotyped trajectories, called manifolds, have previously been reported for *C. elegans* in a neural state-space defined by the first three principal components of the temporal derivative of neural activity (***Kato et al., 2015***). We therefore performed Principal Components Analysis (PCA) on the neural activity (or its temporal derivative) of our recording during the immobilization period, so as to generate a series of principal components or PC’s that capture the major orthogonal components of the variance during immobilization. Population activity during the entire recording was then projected into these first three PCs defined during immobilization, ***Figure 7***c. Neural state space trajectories during immobilization were more structured and stereotyped than during movement and exhibited similarities to previous reports, see ***Figure 7***c,d. Recordings from a second animal was similar and showed pronounced cyclic activity in the first PC of the temporal derivative of neural activity, see ***Figure 7-Figure Supplement 1***b,c. Neural state space trajectories were even more striking and periodic in recordings where the animal had been immobilized for many minutes prior to recording (see ***Figure 7-Figure Supplement 2***, especially PC1). The emergence of structured neural state-space dynamics during immobilization is consistent with the significant change to the correlation structure observed in neural activity. Taken together, these measurements suggest that immobilization alters the correlation structure and dynamics of neural activity.

#### Some neurons change the sign of their correlation with AVA

We further investigated the activity of neuron pair AVA and its correlation to other neurons during movement and immobilization in the recording shown in ***Figure 7***. AVA’s activity was consistent with prior reports. During movement AVA exhibited a sharp rise in response to the animal’s backward locomotion, as expected, see ***Figure 7***b. During immobilization, AVA exhibited slow cycles of activity captured in one of the first three PCs. (AVAL and AVAR received the two largest amplitude weights of all neurons in PC3 dF/dt.) And during both movement and immobilization AVAL and AVAR were consistently highly correlated with one another (*ρ >* 0.89) and participated in a small cluster of positively correlated neurons (most clearly visible in ***Figure 7***e bottom row, small block around AVA).

Interestingly, immobilization induced many neurons to change the sign of their correlations with AVA. For example, some neuron, such as #38 and #0, that had previously been anti-correlated became positively correlated to AVA (Fig ***Figure 8***a,b,d). On average, neurons in this recording become significantly more positively correlated to AVA upon immobilization than during movement (*p* = 0.028 and *p* = 0.038 Welch’s t-test for AVAL and AVAR respectively), ***Figure 8***c.

This suggests that many neurons that appear co-active with AVA during immobilization may not be co-active during movement. Conversely, some neurons that were highly correlated with AVA during movement became anti-correlated during immobilization, such as neurons #71 and #27.

Taken together, our measurements show that immobilization significantly alters the correlation structure of neural activity. Immobilization also causes neurons to change their correlation with known well-characterized neurons, like AVA, from anti-correlated to correlated, or vice versa.

## Discussion

Population activity is important for decoding locomotion in *C. elegans*. Neurons within the population exhibit a diversity of tuning to velocity and curvature and a model that incorporates this diversity outperforms the most predictive single neuron at decoding locomotion. By inspecting the neural weights assigned by our model, we found that only a fraction of neurons are necessary for the model to achieve 90% of its performance. Largely non-overlapping sub-populations of neurons contribute the majority of information for decoding velocity and curvature, respectively.

We found at least two types of neural signals that our model weighted highly for decoding locomotion. One type were neural signals that were strongly correlated with locomotion throughout the duration of the recording. But another type were neural signals with pronounced activity relevant only for particular instances of behavior motifs, for example one deep ventral bend, but not another. By summing up contributions from multiple neurons of this type, the population model was able to capture relevant activity from different neurons at different times to decode all instances of the behavioral motif.

One possible explanation is that superficially similar behavioral features like turns may actually consist of different underlying behaviors. Or the neural representation associated with a turn may also depend on an unobserved behavioral context. The granularity with which to classify behaviors and how to take into account context and behavioral hierarchies remains an active area of research in *C. elegans* (***Broekmans et al., 2016***; ***Liu et al., 2018***; ***Kaplan et al., 2020***) and more broadly (***Berman et al., 2016***; ***Datta et al., 2019***). Ultimately, finding distinct neural signals may help inform our understanding of distinct behavior states and vice versa.

Another possibility is that the same behavior motifs are initiated in the head through different neural pathways. Previous work has suggested that activity in either of two different sets of head interneurons, AVA/AVE/AVD or AIB/RIM, are capable of inducing reversal behavior independently (***Piggott et al., 2011***). If these neurons were active only for the reversals they induced, it could explain why some neurons seem to have activity relevant for some behavioral instances but not others. AVA does not fit this pattern because in our measurements it shows expected activity transients for all reversals. But it is possible that other neurons in the two subsets, or indeed other subsets of neurons, are providing relevant activity for only some instances of a behavior.

Similarly, different sensory modalities such as mechanosensation (***Chalfie et al., 1985***), thermosensation (***Croll, 1975***) and chemosensation (***Ward, 1973***) are known to evoke common behavioral outputs through different sensory neural pathways. Its possible that the neural activities we observe for different behavioral motifs reflects sensory signals that arrive through different sensory pathways to evoke a common downstream motor response.

We specifically investigated the activity and tuning of neuron pair AVAL and AVAR within our population recordings. During simultaneous whole-brain imaging, AVAL and AVAR had activity that was highly correlated with one another, and exhibited the expected peaks in their activity timed to backward locomotion. The temporal derivative of AVA’s activity was correlated to velocity, as expected, but AVA was not in general the most highly correlated neuron to velocity nor was it the neuron most highly weighted by the velocity decoder. This finding highlights the importance of taking a population-based approach at studying neural coding of locomotion.

We found that a linear combination of neural activity and its temporal derivative was suffcient to decode the animal’s locomotion with good performance. Prior studies had investigated neural dynamics of immobilized animals and suggested that particular stereotyped trajectories in neural state space may directly correspond to global motor commands (***Kato et al., 2015***), similar in principle to dynamics in motor cortex during primate reach tasks (***Churchland et al., 2012***). Our measurements suggest that immobilization induces significant changes to the correlation structure of neural dynamics. For example, some neurons altered their correlation with AVA so that a neuron positively correlated with AVA during immobilization was not necessarily positively correlated during movement. Our measurements suggest that neural dynamics from immobilized animals may not entirely reflect the neural dynamics of locomotion.

## Methods

### Strains

Three strains were used in this study, see ***Table 2***. AML32 (***Nguyen et al., 2017***) and AML310 were used for calcium imaging. AML18 (***Nguyen et al., 2016***) served as a calcium insensitive control. Strain AML310 is similar to AML32 but includes additional labels to identify AVA neurons. AML310 was generated by injecting 30 ng/µl of P*rig-3*::tagBFP plasmid into AML32 strains (wtfIs5[P*rab-3*::NLS::GCaMP P*rab-3*::NLS::tagRFP]). AML310 worms were selected and maintained by picking individuals expressing BFP fluorescence in the the head. Animals were cultivated in the dark on NGM plates with a bacterial lawn of OP50.

**Table 2.** Strains used.

| Strain | Genotype | Expression | Role | Reference |
| --- | --- | --- | --- | --- |
| AML310 | wtfIs5[*Prab-3::NLS::GCaMP6s*; *Prab-3::NLS::tagRFP*]; wtfEx258 [*Prig-3::tagBFP::unc-54*] | tag-RFP and GCaMP6s in neuronal nuclei; BFP in cytoplasm of AVA and some pharyngeal neurons (likely I1, I4, M4 and NSM) | Calcium imaging with AVA label | This Study |
| AML32 | wtfIs5[*Prab-3::NLS::GCaMP6s*; *Prab-3::NLS::tagRFP*] | tag-RFP and GCaMP6s in neuronal nuclei | Calcium imaging | (*Nguyen et al., 2017*) |
| AML18 | wtfIs3[*Prab-3::NLS::GFP*, *Prab-3::NLS::tagRFP*] | tag-RFP and GFP in neuronal nuclei | Control | (*Nguyen et al., 2016*) |

### Whole brain imaging

#### Whole brain imaging in moving animals

Whole brain imaging of freely moving animals was performed as described previously (***Nguyen et al., 2016, 2017***). ***Table 3*** lists all recording used in the study and ***Table 4*** cross-lists the recordings according to figure. Briefly, adult animals were placed on an imaging plate (a modified NGM media lacking cholesterol and with agarose in place of agar) and covered with mineral oil. A coverslip was placed on top of the plate with 100 µm plastic spacers between the coverglass and plate surface. The coverslip was fixed to the agarose plate with valap. Animals were recorded on a custom whole brain imaging system, which simultaneously records four video streams to image the calcium activity of the brain while simultaneously capturing the animal’s behavior. We record 10x magnification darkfield images of the body posture, 10x fluorescence images of the head for real-time tracking, and two 40x image streams of the neurons in the head, one showing tagRFP and one showing either GCaMP6s, GFP, or BFP. The 10x images are recorded at 50 frames/s, and the 40x fluorescence images are recorded at a rate of 200 optical slices/s, with a resulting acquisition rate of 6 head volumes/s. Recordings were stopped when the animal ran to the edge of the plate,when they left the field of view, or when photobleaching decreased the contrast between tag-RFP and background below a minimum level. Intensity of excitation light for fluorescent imaging was adjusted from recording to recording to achieve different tradeoffs between fluorescence intensity and recording duration.

**Table 3.**
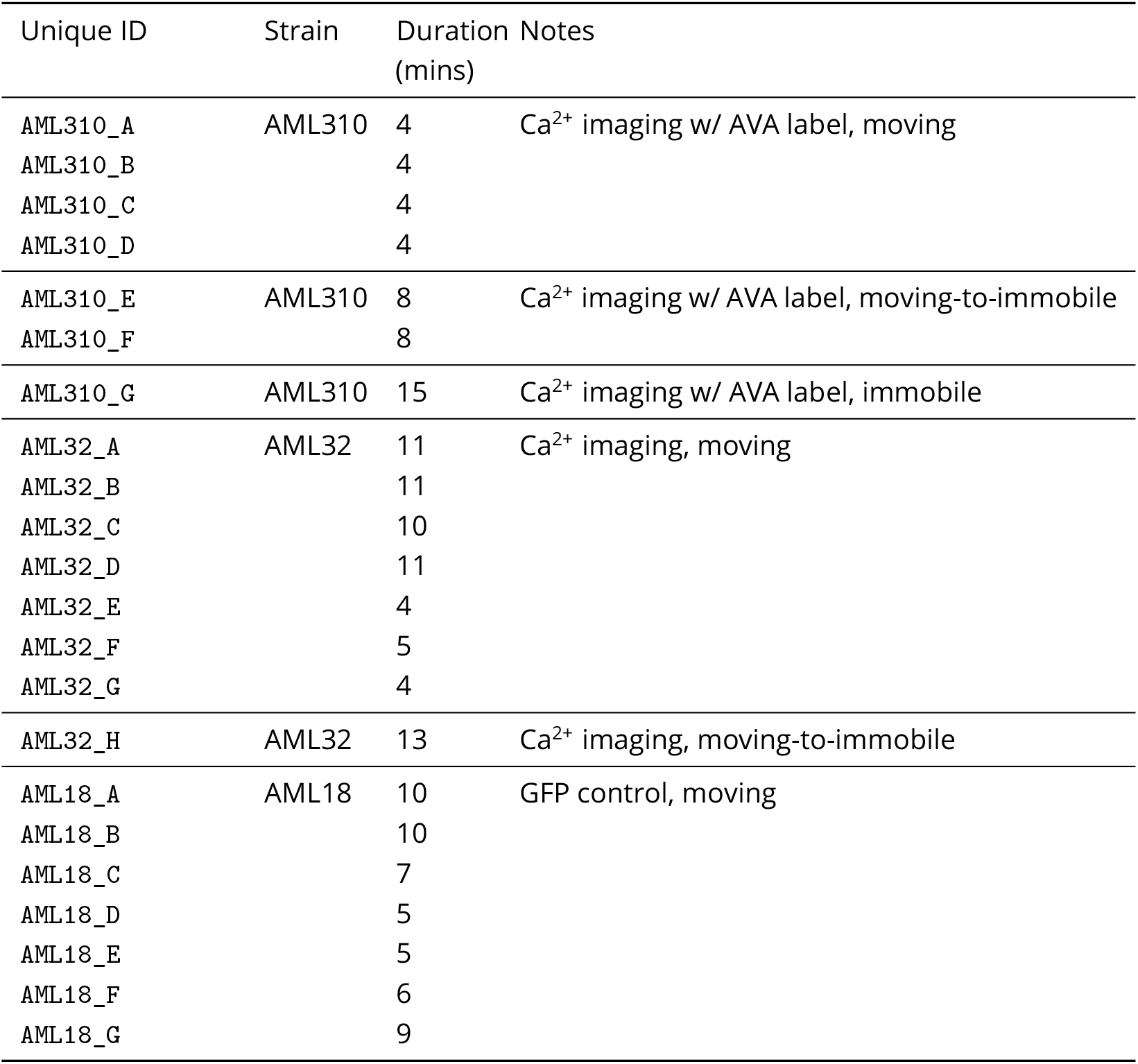
Recordings used in this study.

**Table 4.** List of recordings included in each figure

| Figure | Recordings |
| --- | --- |
| **Figure 1; Figure 1 - Figure Supplement 1; Figure 2a,b** | AML310_A |
| **Figure 2c** | AML310_A-D |
| **Figure 3a-d** | AML310_A |
| **Figure 3e,f; Figure 3-Figure Supplement 2** | AML310_A-D, AML32_A-G, AML18_A-G |
| **Figure 3 - Figure Supplement 1; Figure 3 - Figure Supplement 3** | AML310_A-D, AML32_A-G |
| **Figure 4** | AML32_A |
| **Figure 5; Figure 5 - Figure Supplement 1; Figure 5 - Figure Supplement 2; Figure 6a,b** | AML310_A |
| **Figure 6c** | AML310_A-D, AML32_A-G |
| **Figure 7a-f** | AML310_E |
| **Figure 7g** | AML310_A-F, AML32_A-H |
| **Figure 7 - Figure Supplement 1** | AML32_H |
| **Figure 7 - Figure Supplement 2** | AML310_G |
| **Figure 8** | AML310_E |

Freely moving recordings had to meet the following criteria. The animal had to be active and the recording had to be at least 200 seconds. The tag-RFP neurons also had to be successfully segmented and tracked via our analysis pipeline.

#### Moving to immobile transition experiments

Adult animals were placed in a PDMS microfludic artificial dirt style chip (***Lockery et al., 2008***) filled with M9 medium where the animal could crawl. The chip was imaged on the whole brain imaging system. A computer controlled microfluidic pump system delivered either M9 buffer or M9 buffer with the paralytic levamisole or tetramisole to the microfluidic chip. Calcium activity was recorded from the worm as M9 buffer flowed through the chip with a flow rate of order a milliliter a minute. Partway through the recording, the drug buffer mixture was delivered at the same flow rate. At the conclusion of the experiment for AML310 worms, BFP was imaged.

Different drug concentrations were tried for different recordings to find a good balance between rapidly immobilizing the animal without also inducing the animal to contract and deform. Paralytic concentrations used were: 400 *µ*M for AML310_E, 100 *µ*M for AML310_F, and 5 *µ*M for AML32_H.

Recordings were performed until a recording achieved the following criteria for inclusion: 1) the animal showed robust locomotion during the moving portion of the recording, including multiple reversals. 2) The animal quickly immobilized upon application of the drug. 3) The animal remained immobilized for the remainder of the recording except for occasional twitches, 4) the immobilization portion of the recording was of sufficient duration to allow us to see multiple cycles of the stereotyped neural state space trajectories if present and 5) for strain AML310, neurons AVAL and AVAR were required to be visible and tracked throughout the entirety of the recording. For the statistics of correlation structure in ***Figure 7***f, recording AML310_F was also included even though it did not meet all criteria (it lacked obvious reversals).

#### Whole brain imaging in immobile animals

We performed whole brain imaging in adult animals immobilized with 100 nm polystyrene beads (***Kim et al., 2013***). The worms were then covered with a glass slide, sealed with valap, and imaged using the Whole Brain Imager.

#### Neuron segmentation, tracking and fluorescence extraction

Neurons were segmented and tracked using the the Neuron Registration Vector Encoding (NeRVE) and clustering approach described previously (***Nguyen et al., 2017***) with minor modifications which are highlighted below. As before, video streams were spatially aligned with beads and then synchronized using light flashes. The animals’ posture was extracted using an active contour fit to the 10x darkfield images. But in a departure from the method in (***Nguyen et al., 2017***), the high magnification fluorescent images are now straightened using a different centerline extracted directly from the fluorescent images. As in (***Nguyen et al., 2017***), the neural dynamics were then extracted by segmenting the neuronal nuclei in the red channel and straightening the image according to the body posture. Using repeated clustering, neurons are assigned identities over time. The GCaMP signal was extracted using the neural positions found from tracking. The pipeline returns datasets containing RFP and GCaMP6s fluorescence values for each successfully tracked neuron over time, and the centerline coordinates describing the posture of the animal over time. These are subsequently processed to extract neural activity or behavior features.

The paralytic used in moving-to-immobile recordings (***Figure 7***) caused the animal’s head to contract, which would occasionally confuse our tracking algorithm. In those instances the automated NeRVE tracking and clustering was run separately on the moving and immoibile portions of the recording (before and after contraction), and then a human manually tracked neurons during the transition period (one to two minutes) so as to stitch the moving and immobile tracks together.

#### Photobleaching correction, outlier detection and pre-processing

The raw extracted RFP or GCaMP fluorescent intensity timeseries were preprocessed to correct for photobleaching. Each time-series was fit to a decaying exponential. Those that were well fit by the exponential were normalized by the exponential and then rescaled to preserve the timeseries’ original mean and variance as in (***Chen et al., 2019***). Timeseries that were poorly fit by an exponential were left as is. If the majority of neurons in a recording were poorly fit by an exponential, this indicated that the animal may have photobleached prior to the recording and the recording was discarded.

Outlier detection was performed to remove transient artifacts from the fluorescent time series. Fluorescent time points were flagged as outliers and omitted if they met any of the following conditions: the fluorescence deviated from the mean by a certain number of standard deviations (*F <* −2*σ*, or *F >* 5*σ*, for RFP; *F >* 5*σ*, for GCaMP); the RFP fluorescence dropped below a threshold; the ratio of GCaMP to RFP fluorescence dropped below a threshold; a fluorescence timepoint was both preceded by and succeeded by missing timepoints or values deemed to be outliers; or if the majority of other neurons measured during the same volume were also deemed to be outliers.

Fluorescent time series were smoothed by convolution with a Gaussian (*σ*, = 0.83s) after interpolation. Omitted time points, or gaps where the neuron was not tracked, were excluded from single-neuron analyses, such as the calculation of each neuron’s tuning curve. It was not practical to exclude missing time points from population-level analyses such as linear decoding or Principal Components Analysis. In these population-level analyses, interpolated values were used. Time points in which the majority of neurons has missing fluoresecent values were excluded, even in population level analyses. Those instances are shown as white vertical stripes in the fluorescent activity heat maps shown in ***Figure 1*** and ***Figure 3-Figure Supplement 2***.

#### Motion-correction

We used the GCaMP fluorescence together with the RFP fluorescence to calculate a motion corrected fluorescence, *F*_*mc*_ used through the paper. Note sometimes the subscript _mc_ is omitted for brevity. Motion and deformation in the animal’s head can introduce artifacts into the fluorescent time-series. Many of these artifacts should be common to both GCaMP and RFP fluorescence. For example, if a neuron is compressed during a head bend, the density of both GCaMP and RFP should increase, causing an increase in the fluorescence in both time-series. Moreover, the RFP time series should be dominated by artifacts because, in the absence of motion, the RFP fluorescent intensity would be constant. We therefore sought to reject fluctuations common to both GCaMP and RFP time series by using independent components analysis (ICA). ICA has previously been used in neuroscience to identify spikes in intracranial recordings (***Kobayashi et al., 2001***) or to automatically define regions of interest from large-scale calcium recordings (***Mukamel et al., 2009***). Here we use ICA to find a motion corrected fluorescent signal, *F*_mc_, that captures the activity in the GCaMP timeseries *F*_GCaMP_ but rejects fluctuations common to both the GCaMP and RFP signals, *F*_RFP_ and *F*_RFP_. We performed ICA on the normalized mean-subtracted time series and extracted two components (***Pedregosa et al., 2011***). The component most correlated with *F*_RFP_ was deemed noise and rejected. The remaining component was deemed the motion corrected fluorescence signal *F*_mc_ and was re-scaled and offset to retain the mean, variance and sign of the original *F*_GCaMP_ time series.

#### Temporal derivative

The temporal derivatives of motion corrected neuron signals are estimated using a Gaussian derivative kernel of width 2 3 s. For brevity we denote this kernel-based estimate as 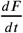.

#### Identifying AVA

AVAL and AVAR were identified in recordings of AML310 by their known location and the presence of a BFP fluorescent label expressed under the control of *rig-3* promoter. BFP was imaged immediately after calcium imaging was completed, usually while the worm was still moving freely. To image BFP, a 488 nm laser was blocked and the worm was then illuminated with 405 nm laser light. In one of the recordings, only one of the two AVA neurons was clearly identifiable throughout the duration of the recording. For that recording, only one of the AVA neurons was included in analysis.

### Measuring and representing locomotion

To measure the animal’s velocity and body curvature we used an analysis based on an eigenvalue decomposition of posture (***Stephens et al., 2008***), as follows. Centerlines of the worms were extracted from the whole-brain imaging recordings using an active contour fit to a darkfield image of the worm as in (***Nguyen et al., 2017***). The resulting centerlines were projected onto a 4-dimensional basis set of pose eigenvectors that had previously been computed from an eigenvalue analysis of centerlines of 135,958 frames of freely moving worms taken from an in-house collection of recordings on the whole brain imager. The first four eigenvectors explained 96% of the observed variance of the in-house collection of recordings.

The projection of the centerlines onto the eigenvectors results in a timeseries of coeffcients, one per eigenvector. Two of the coefficients describe the body bends that the worm creates during its sinusoidal locomotion. The phase of the body bends is 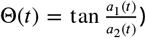, where *a*_1_ and *a*_2_ are the coeffcients for the first two eigenvectors. The derivative of the phase *d*Θ/*dt* is the phase velocity that describes the speed of bend propagation in the worm. We chose to use this velocity because it more directly reflects the animal’s muscle output than the animal’s center of mass velocity, which also involves mechanical interactions with the substrate and is affected, for example, by slip. Per our convention *d*0/*dt* is reported in radians per second and is positive for forward motion and negative during a reversal. Velocity is obtained by filtering 0 with a gaussian derivative filter with width of *σ*, = 2s. The third coeffcient *a*_3_(*t*) is a dimensionless quantity that corresponds to the body curvature of the animal, which is related to turning. For example, when the animal initiates a turn its velocity is positive, and its body curvature is large. The sign of *a*_3_(*t*) describes the dorsal-ventral bend direction. *a*_3_(*t*) is referred to as curvature throughout this work.

### Relating neural activity to behavior

#### Tuning Curves

The Pearson’s correlation coeffcient *ρ* is reported for each neurons’ tuning, as in ***Figure 1***d,e. To reject the null hypothesis that a neuron is correlated with behavior by chance we took a shuffing approach and applied a Bonforroni correction for multiple hypothesis testing. We shuffed our data in such a way as to preserve the correlation structure in our recording. To calculate the shuffe, each neuron’s activity was time-reversed and circularly shifted relative to behavior by a random time lag and then the Pearson’s correlation coeffcient was computed. Shuffing was repeated for each neuron in a recording 5,000 times to build up a distribution of 5000*N* values of *ρ*, where *N* is the number of neurons in the recording. To reject the null hypothesis at 0.05% confidence, we apply a Bonforonni correction such that a correlation coeffcient greater than *ρ* (or less than, depending on the sign) must have been observed in the shuffled distribution with a probability less than 0.05/(2*N*). The factor of 2*N* arises from accounting for multiple hypothesis testing for tuning of both *F* and *dF* /*dt* for each neuron.

#### Population Model

We use a ridge regression (***Hoerl and Kennard, 1970***) model to decode behavior signals *y*(*t*) (the velocity and the body curvature). The model prediction is given by a linear combination of neural activities and their time derivatives,

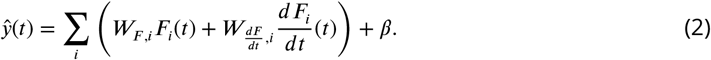

Note here we are omitting the _mc_ subscript for convenience, but these still refer to the motion corrected fluorescence signal.

We scale all these features to have zero mean and unit variance, so that the magnitudes of weights can be compared to each other. To determine the parameters 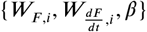 we hold out a test set comprising the middle 40% of the recording, and use the remainder o f the data for training. We minimize the cost function

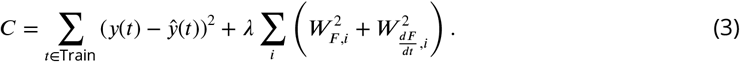

The hyperparameter *λ*sets the strength of the ridge penalty in the second term. We choose *λ*by splitting the training set further into a second training set and a cross-validation set, and training on the second training set with various values of λ. We choose the value which gives the best performance on the cross-validation set.

To evaluate the performance of our model, we use a mean-subtracted coeffcient of determination metric, 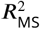, on the test set. This is defined by

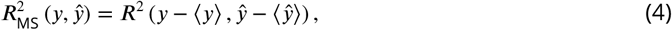

where we use the conventional definition of *R*^2^, defined here for an arbitrary true signal *z* and corresponding model prediction 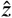:

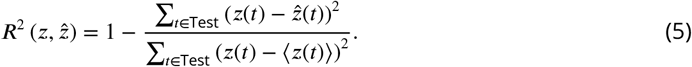

Note that 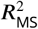can take any value on (−∞, 1].

#### Restricted models

To assess the distribution of locomotive information throughout the animal’s brain, we compare with two types of restricted models. First, we use a Best Single Neuron model in which all but one of the coeffcients 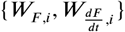 in (2) are constrained to vanish. We thus attempt to represent behavior as a linear function of a single neural activity, or its time derivative These models are shown in ***Figure 3***. Second, after training the population model, we sort the neurons in descending order of 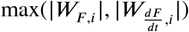. We then construct models using a subset of the most highly weighted neurons, with the relative weights on their activities and time derivatives fixed by those used in the population model. The performance of these truncated models can be tabulated as a function of the number of neurons included to first achieve a given performance, as shown in ***Figure 6***.

#### Alternative models

The population model used throughout this work refers to a linear model with derivatives using ridge regression. In ***Figure 3 - Figure Supplement 3***, we show the performance of seven alternative population models at decoding velocity for our exemplar recording. The models are summarized in ***Table 5***. Many of these models perform roughly as well as the linear population model used throughout the paper. Our chosen model was selected both for its relative simplicity and because it showed the highest mean performance at decoding velocity across recordings.

**Table 5.** Alternative models explored. Most are linear models, using either the Ridge or ElasticNet regularization. In some cases we add an additional term to the cost function which penalizes errors in the temporal derivative of model output (which, for velocity models, corresponds to the error in the predicted acceleration). For features, we use either the neural activities alone, or the neural activities together with their temporal derivatives. We also explore two nonlinear models: MARS ***Friedman*** (***1991***), and a shallow decision tree which chooses between two linear models.

| Model | Penalty | Features | Number of Parameters |
| --- | --- | --- | --- |
| Linear | Ridge | $F$ and $dF/dt$ | $2N_n + 1$ |
| Linear | Ridge | $F$ | $N_n + 1$ |
| Linear | Ridge + Acceleration Penalty | $F$ and $dF/dt$ | $2N_n + 1$ |
| Linear | Ridge + Acceleration Penalty | $F$ | $N_n + 1$ |
| Linear | ElasticNet | $F$ and $dF/dt$ | $2N_n + 1$ |
| Linear | ElasticNet | $F$ | $N_n + 1$ |
| MARS (nonlinear) | MARS | $F$ and $dF/dt$ | variable |
| Linear with Decision Tree (nonlinear) | Ridge | $F$ and $dF/dt$ | $4N_n + 9$ |

***Figure 3 - Figure Supplement 3***a-b show the model we use throughout the paper, and the same model but with only fluorescence signals (and not their time derivatives) as features. The latter model attains a slightly lower score of 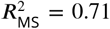. Note that while adding features is guaranteed to improve performance on the training set, performance on the test set did not necessarily have to improve. Nonetheless, we generally found that including the time derivatives led to better predictions on the test set.

***Figure 3 - Figure Supplement 3***c-d show a variant of the linear model where we add an acceleration penalty to the model error. Our cost function becomes (cf. (3))

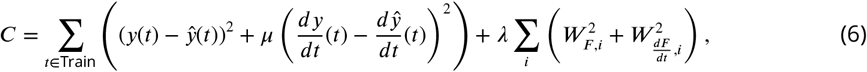

where the derivatives 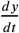 and 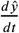 are estimated using a Gaussian derivative filter. The parameter *µ* is set to 10. For our exemplar recording, adding the acceleration penalty hurts the model when derivatives are not included as features, but has little effect when they are.

***Figure 3 - Figure Supplement 3***e-f show a variant where we use an ElasticNet penalty instead of a ridge penalty (***Zou and Hastie, 2005***). If we write the ridge penalty as the *L*_2_ norm of the weight vector, so that

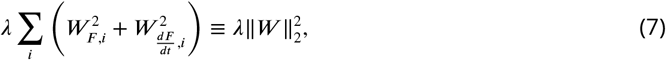

the ElasticNet penalty is defined by

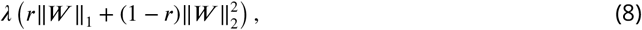

where

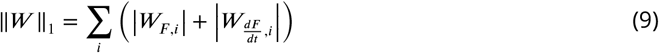

is the *L*_1_ norm of the weight vector. The quantity is known as the *L*_1_ ratio, and in ***Figure 3 - Figure Supplement 3*** it is set to 10^−2^. We have also tried setting via cross-validation, and found similar results.

***Figure 3 - Figure Supplement 3***g uses the multivariate adaptive regression splines (MARS) model (***Friedman, 1991***). The MARS model incorporates nonlinearity by using rectified linear functions of the features, or products of such functions. Generally they have the advantage of being more flexible than linear models while remaining more interpretable than a neural network or other more complicated nonlinear model. However, we find that MARS somewhat underperforms a linear model on our data.

***Figure 3 - Figure Supplement 3***h uses a decision tree classifier trained to separate the data into forward-moving and backward-moving components, and then trains separate linear models on each component. For our exemplar recording, this model performs slightly better than the model we use throughout the paper. This is likely a result of the clear AVAR signal in Figure 2, which can be used by the classifier to find the backward-moving portions of the data. Across all our recordings, this model underperforms the simple linear model.

### Correlation structure analysis

The correlation structure of neural activity was visualised as the correlation matrix, *ρ*_*i*_, *j*. To observe changes in correlation structure, a correlation matrix for the moving portion of the recording was calculated separately from the immobile portion. The time immediately following delivery of the paralytic when the animal was not yet paralzed was excluded (usually one to two minutes). To quantify the magnitude of the change in correlation structure, a dissimilarity metric was defined as the root mean-squared change in each neuron’s pairwise correlations, 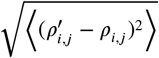. As a control, changes to correlation structure were measured in freely moving animals. In this case the correlation structure of the first 30% of the recording was compared to the correlation structure of latter 60% of the recording, so as to mimic the relative timing in the moving-to-immobile recordings.

## Software

Analysis scripts are available at https://github.com/leiferlab/PredictionCode

## Data

Data from all experiments including calcium activity traces and animal pose and position are publicly available at https://doi.org/10.17605/OSF.IO/R5TB3

## Acknowledgments

This work was supported by grants from the Simons Foundation (SCGB #324285, and SCGB #543003, AML). This work was supported in part by the National Science Foundation, through the Center for the Physics of Biological Function (PHY-1734030) and an NSF CAREER Award to AML (IOS-1845137). ANL is supported by a National Institutes of Health institutional training grant NIH T32 MH065214 through the Princeton Neuroscience Institute. Strains are distributed by the CGC, which is funded by the NIH Offce of Research Infrastructure Programs (P40 OD010440).

## Author contributions

- Kelsey Hallinen: Formal Analysis, Investigation, Visualization, Writing - original draft, Writing - review and editing.
- Ross Dempsey: Formal Analysis, Investigation, Methodology, Software, Visualization, Writing original draft, Writing - review and editing.
- Monika Scholz: Formal Analysis, Investigation, Methodology, Software, Visualization, Writing original draft, Writing - review and editing.
- Xinwei Yu: Formal Analysis, Investigation, Methodology, Resources, Software, Writing - review and editing
- Ashley Linder: Formal Analysis, Investigation, Methodology, Resources, Software, Writing - review and editing
- Francesco Randi: Resources, Writing - review and editing, Developed optical instrument.
- Anuj Sharma: Resouresc, Writing - review and editing, Generated all transgenics.
- Joshua Shaevitz: Conceptualization, Supervision, Funding Acquisition, Writing - review and editing.
- Andrew Leifer: Conceptualization, Formal Analysis, Funding Acquisition, Project administration, Software, Supervision, Visualization, Writing - original draft, Writing - review and editing.

---

**Figure S1–Figure supplement 1.**
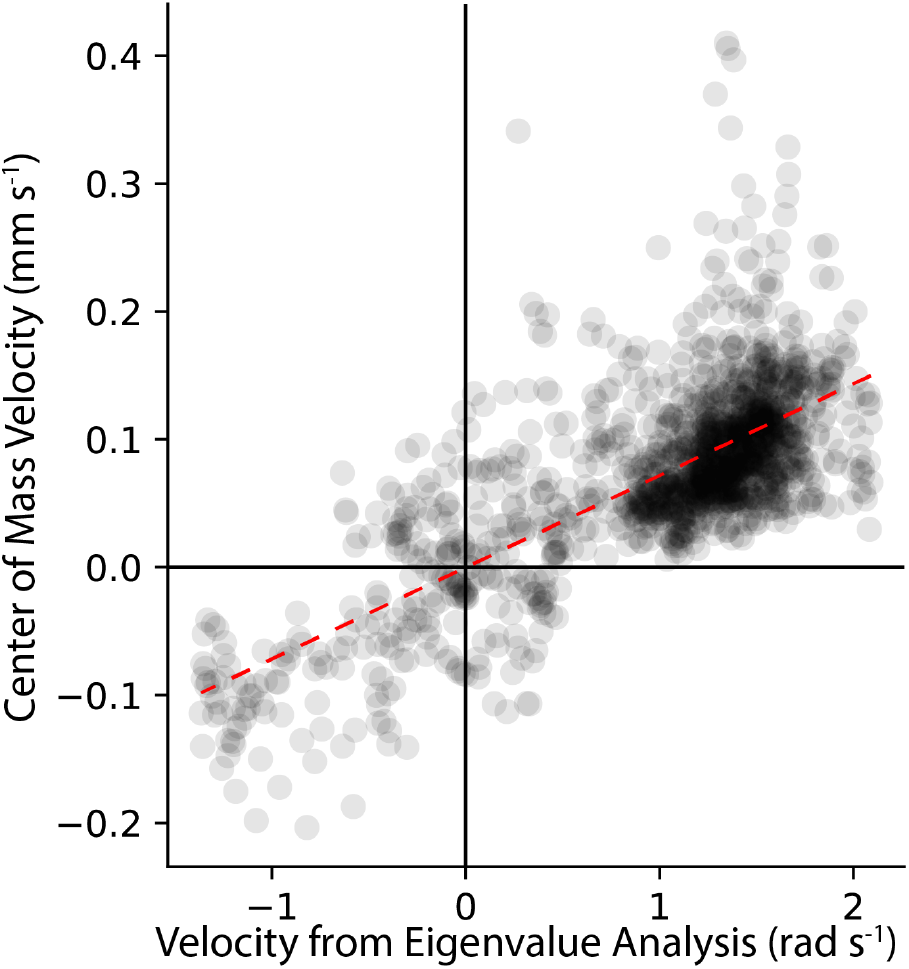
Comparison of center-of-mass velocity and eigenvalue decomposition-derived velocity for recording `AML310_A` . Dashed line is line of best fit.

**Figure S3–Figure supplement 1.**
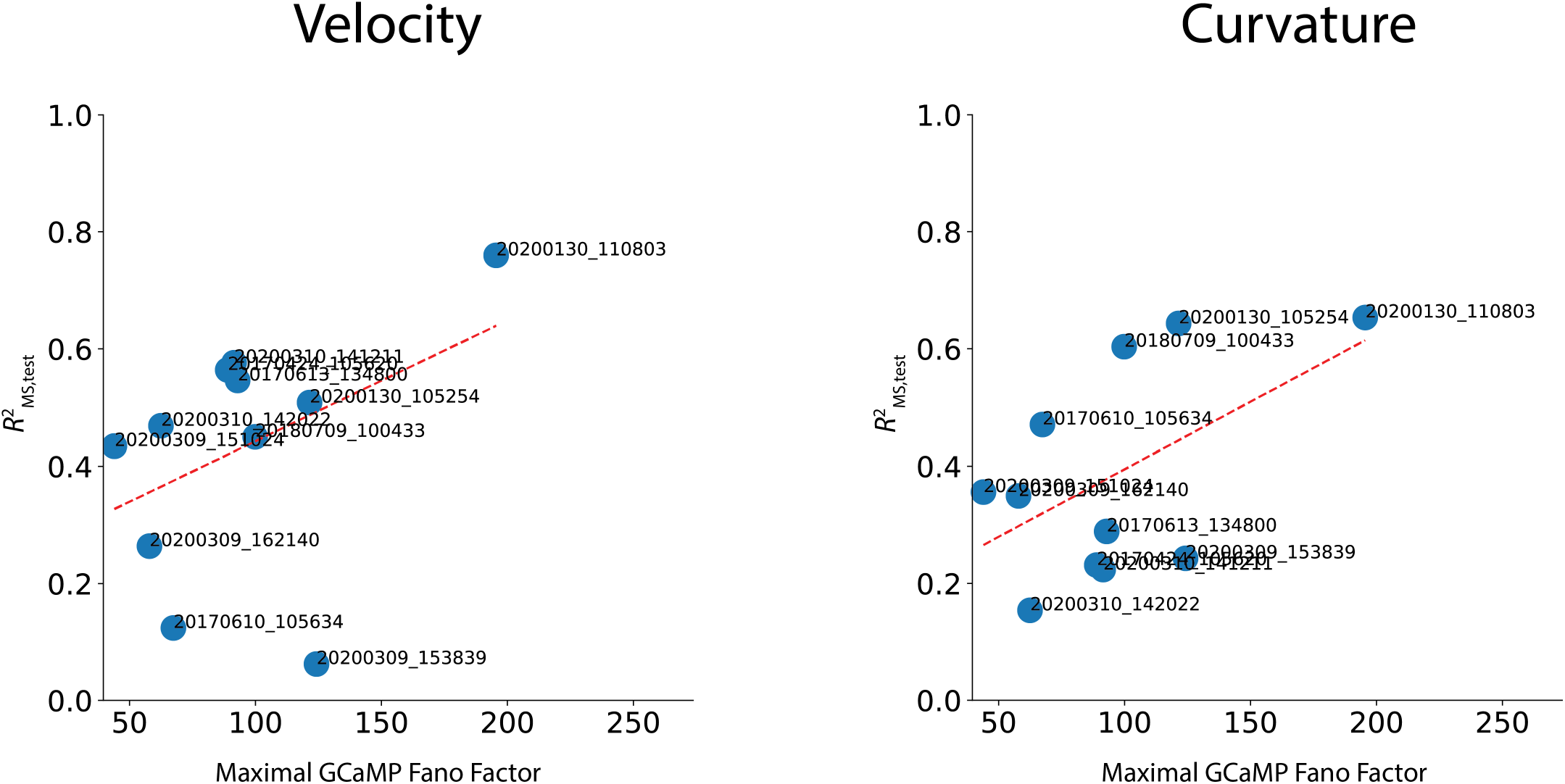
Performance correlates with maximal GCaMP Fano Factor, a metric of signal-to-noise ratio. Decoding performance is plotted against maximal GCaMP Fano Factor for each recording for velocity and curvature. Maximal GCaMP Fano Factor is the Fano Factor of the raw GCaMP activity for the neuron in each recording with the highest Fano Factor, 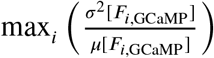 Labels for each recording are shown. Dashed red line is the line of best fit (correlation coefficient between fit and data is *ρ* = 0.42 for velocity and *ρ* = 0.53 for curvature.)

**Figure S3–Figure supplement 2.**
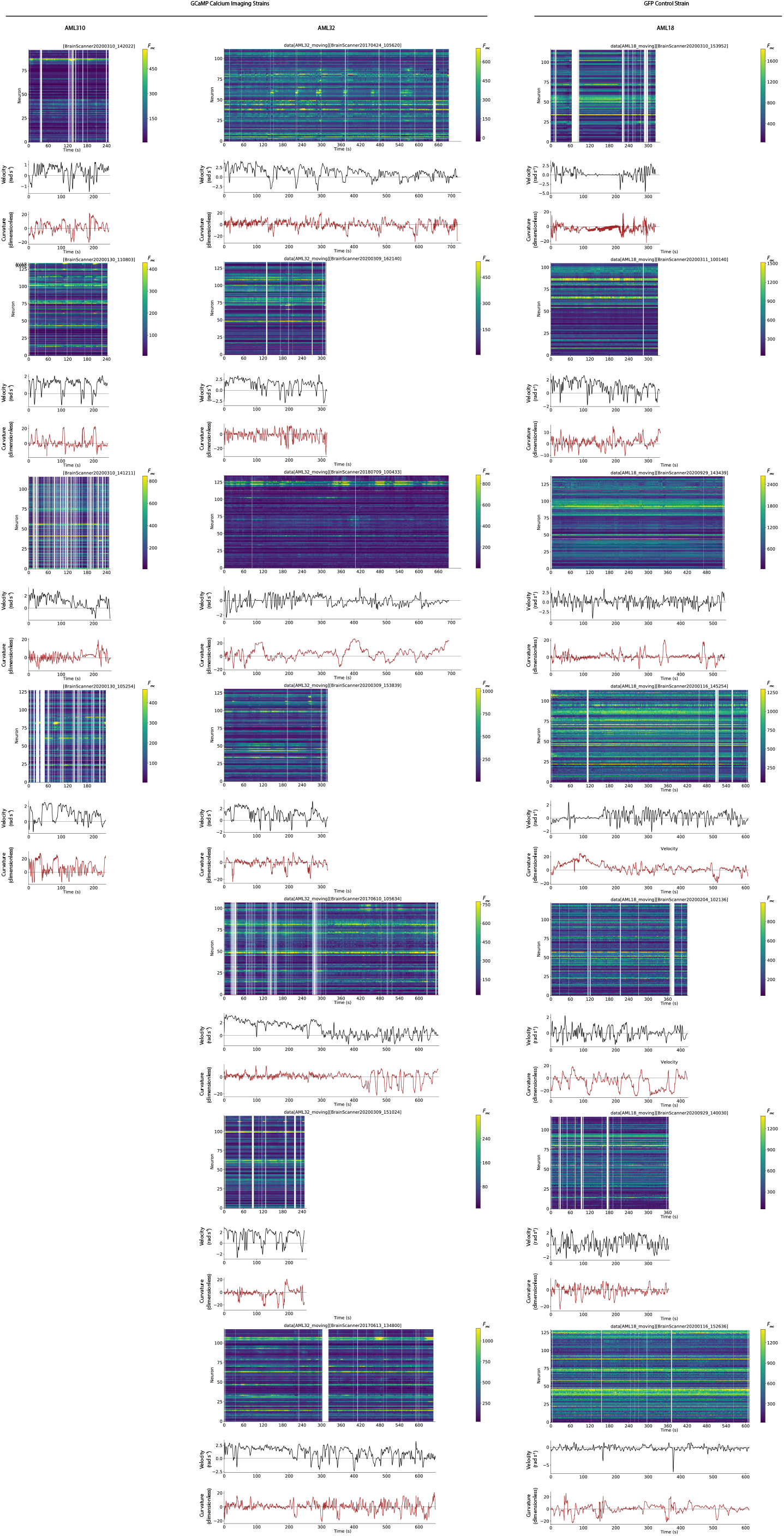
Neural activity and behavior for all freely moving recordings, including GCaMP imaging strains (AML310 and AML32) and GFP control strains (AML18).

**Figure S3–Figure supplement 3.**
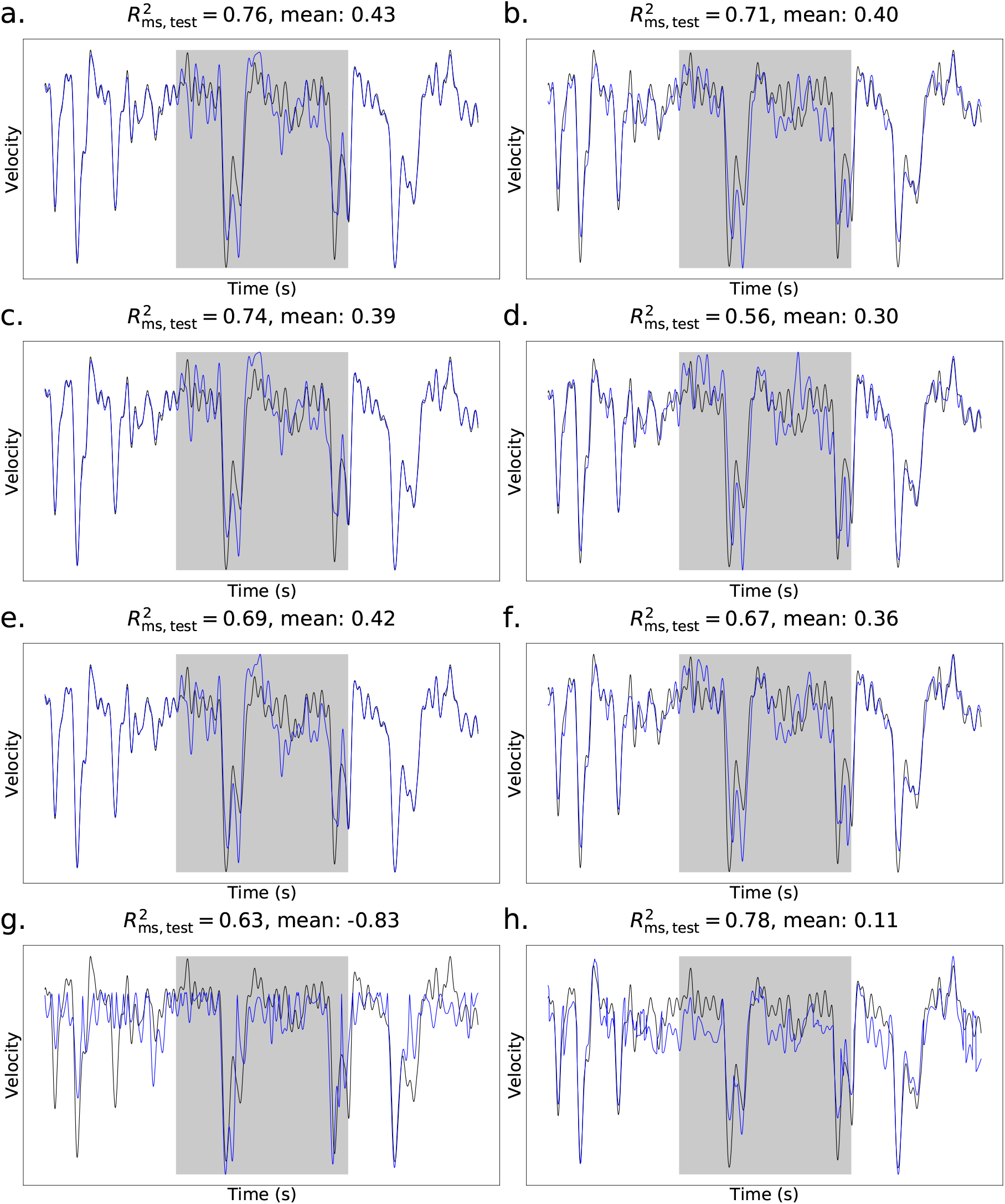
Performance of alternative population models for decoding velocity. Traces are shown for exemplar recording `AML310_A` . Mean across all recordings is also listed. a.) The population model used throughout the paper. This model uses ridge regression with fluorescence signals and their temporal derivatives as features. b.) A linear model using ridge regression, with only fluorescence signals as features. c.) A linear model using fluorescence signals and their temporal derivatives as features, regularized with a combination of a ridge penalty and the squared error of the temporal derivative of behavior. d.) The model in c., but using only fluorescence signals as features. e.) A linear model using fluorescence signals and their temporal derivatives as features, regularized with an ElasticNet penalty with an *L*_1_ ratio of 10^−2^. f.) The model in e., but using only fluorescence signals as features. g.) The multivariate adaptive regression splines (MARS) model, using fluorescence signals and their temporal derivatives as features. h.) A linear model together with a shallow decision tree, using fluorescence signals and their temporal derivative as features.

**Figure S5–Figure supplement 1.**
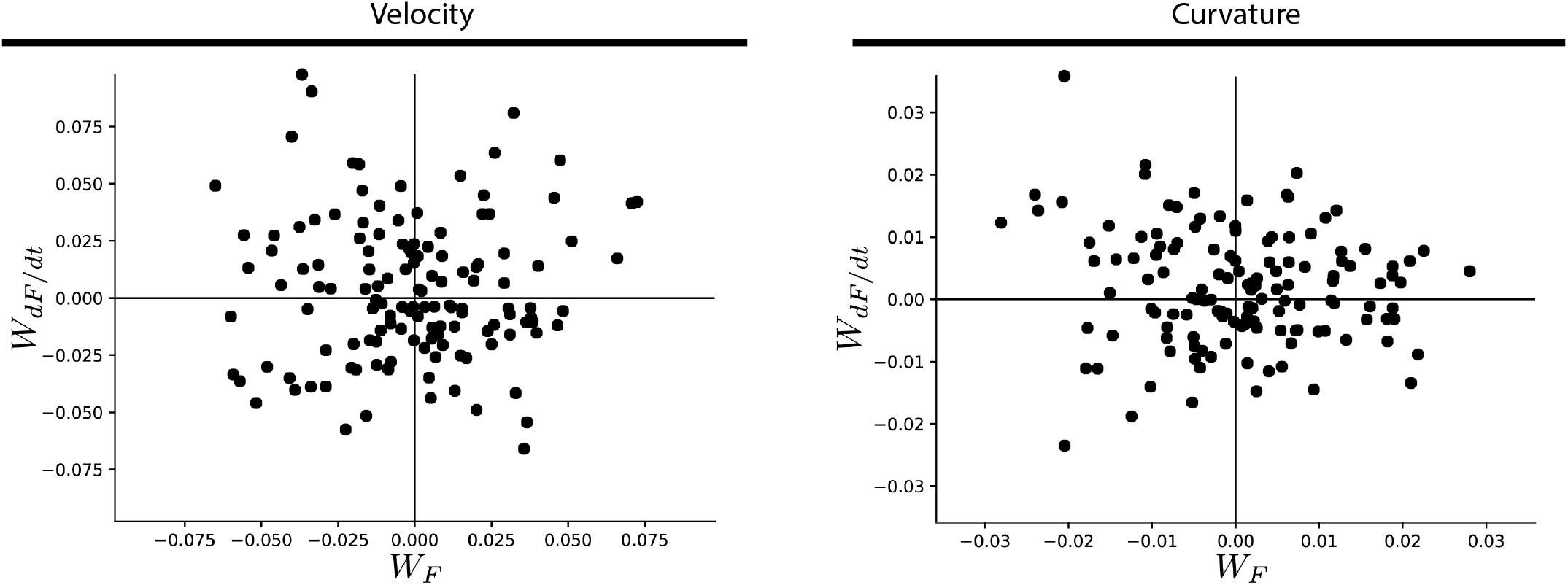
Comparison of weights assigned to a neuron’s activity versus its temporal derivative for velocity (left) or curvature (right) decoders. Comparison of weights assigned to a neuron’s activity *W*_*F*_ by the population decoder, versus the weights assigned to its temporal derivative |*W*_*dF* /*dt*_|for each neuron in recording AML310_A.

**Figure S5–Figure supplement 2.**
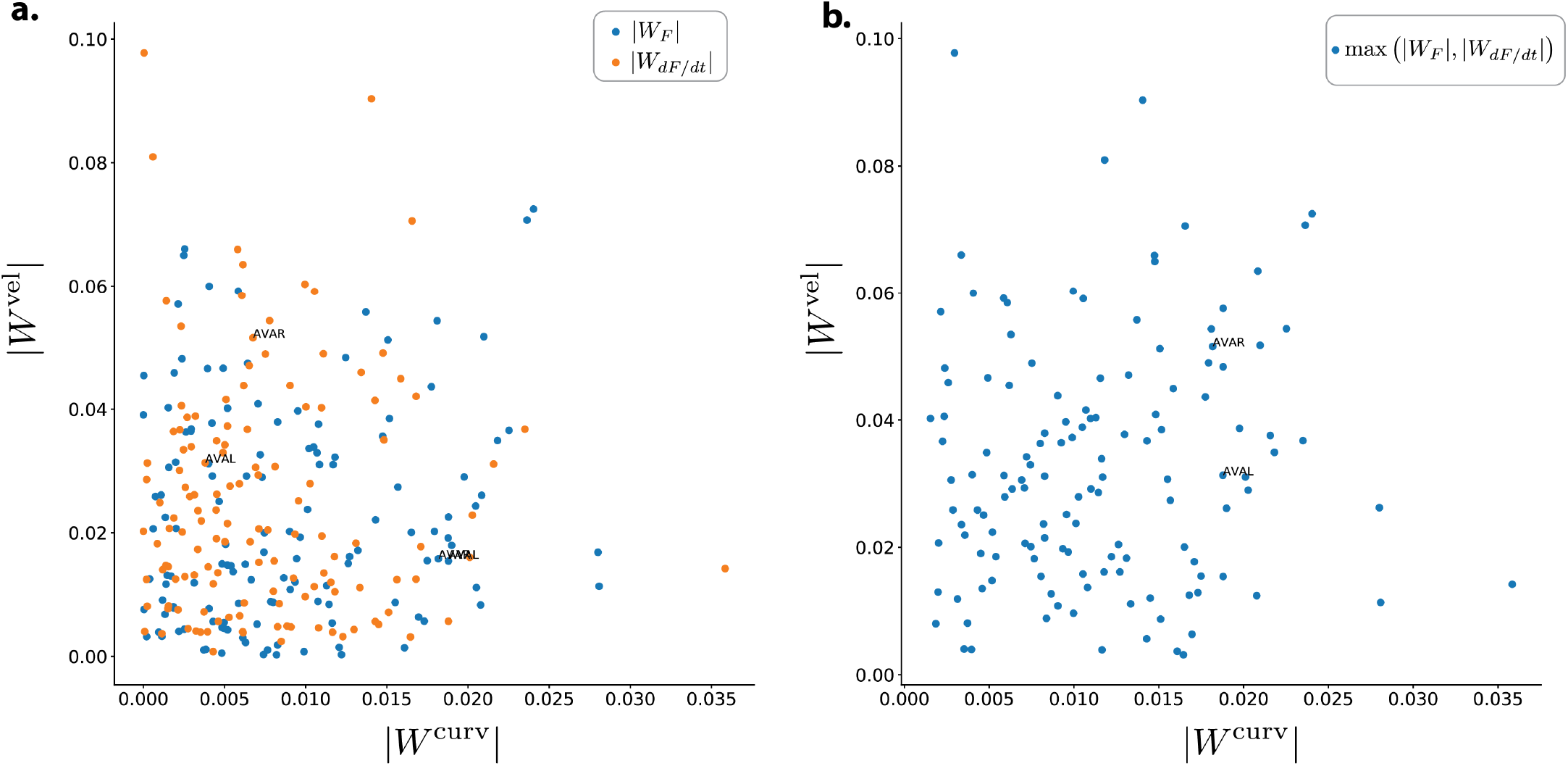
Comparison of weights assigned for decoding velocity vs decoding curvature. a.) The magnitude of the weight assigned to each neuron in recording AML310_A for velocity *W* ^vel^ is compared to the magnitude of its assigned weight for curvature *W* ^curv^ . Each neuron is plotted twice, once for the weight assigned to its activity and once for the weight assigned to the temporal derivative of its activity. b) Same as in (a), except here the higher weight of either activity or its temporal derivative is plotted.

**Figure S7–Figure supplement 1.**
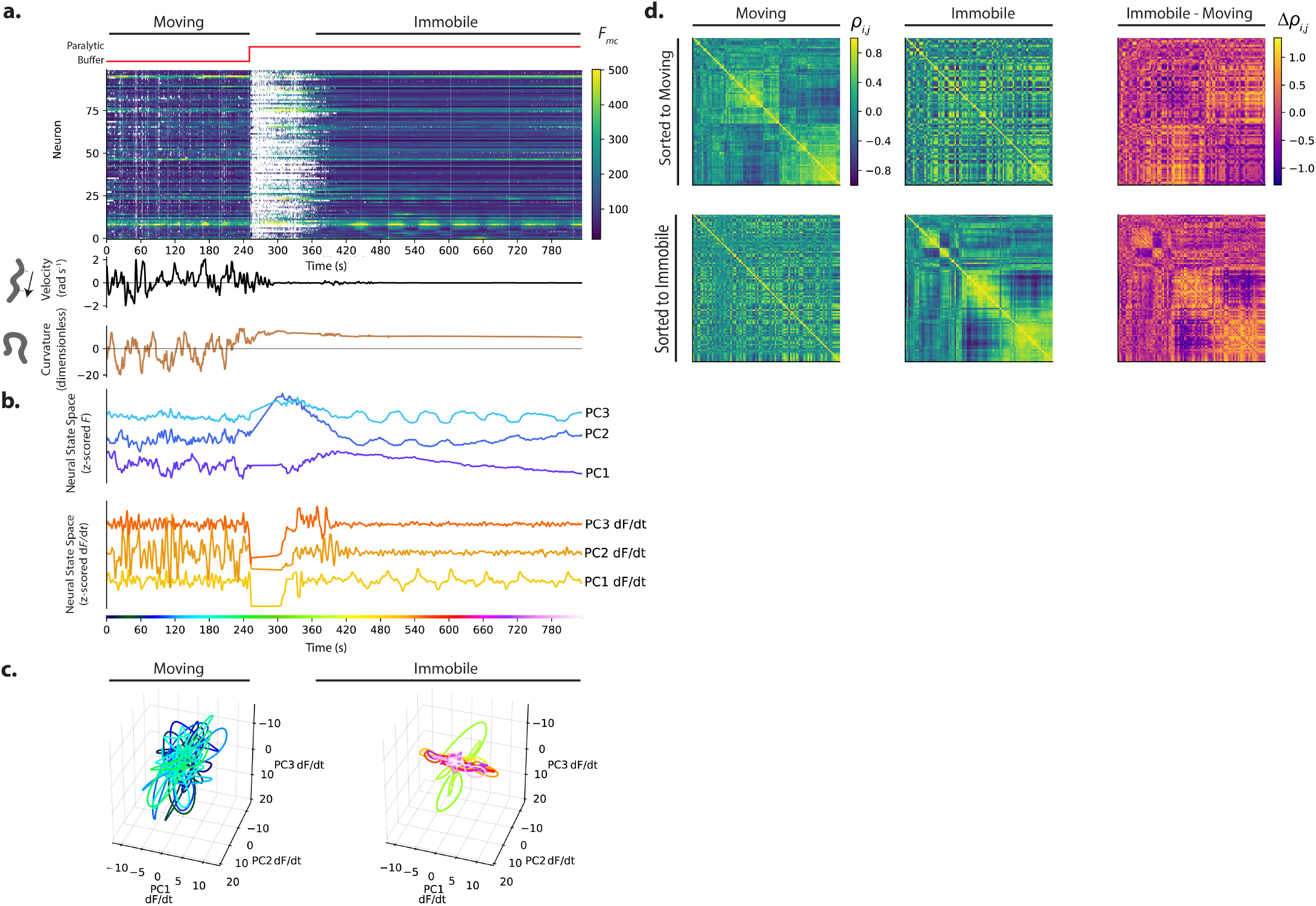
Calcium activity is recorded from an animal as it moves and then is immobilized with a paralytic drug, recording AML32_H . Activity and behavior. b) Population activity (or its derivative) from (a) is shown projected onto its first three PCs, as determined by only the immobilized portion of the recording. c) Neural state space trajectories from (b) are plotted in 3D and shown split into moving and immobile portions. d) Pairwise correlations of neural activity *ρ*_*i,j*_ are shown as heatmaps for all neurons during movement and immobilization, sorted via clustering algorithm. Top and bottom rows are sorted to movement or immobilization, respectively.

**Figure S7–Figure supplement 2.**
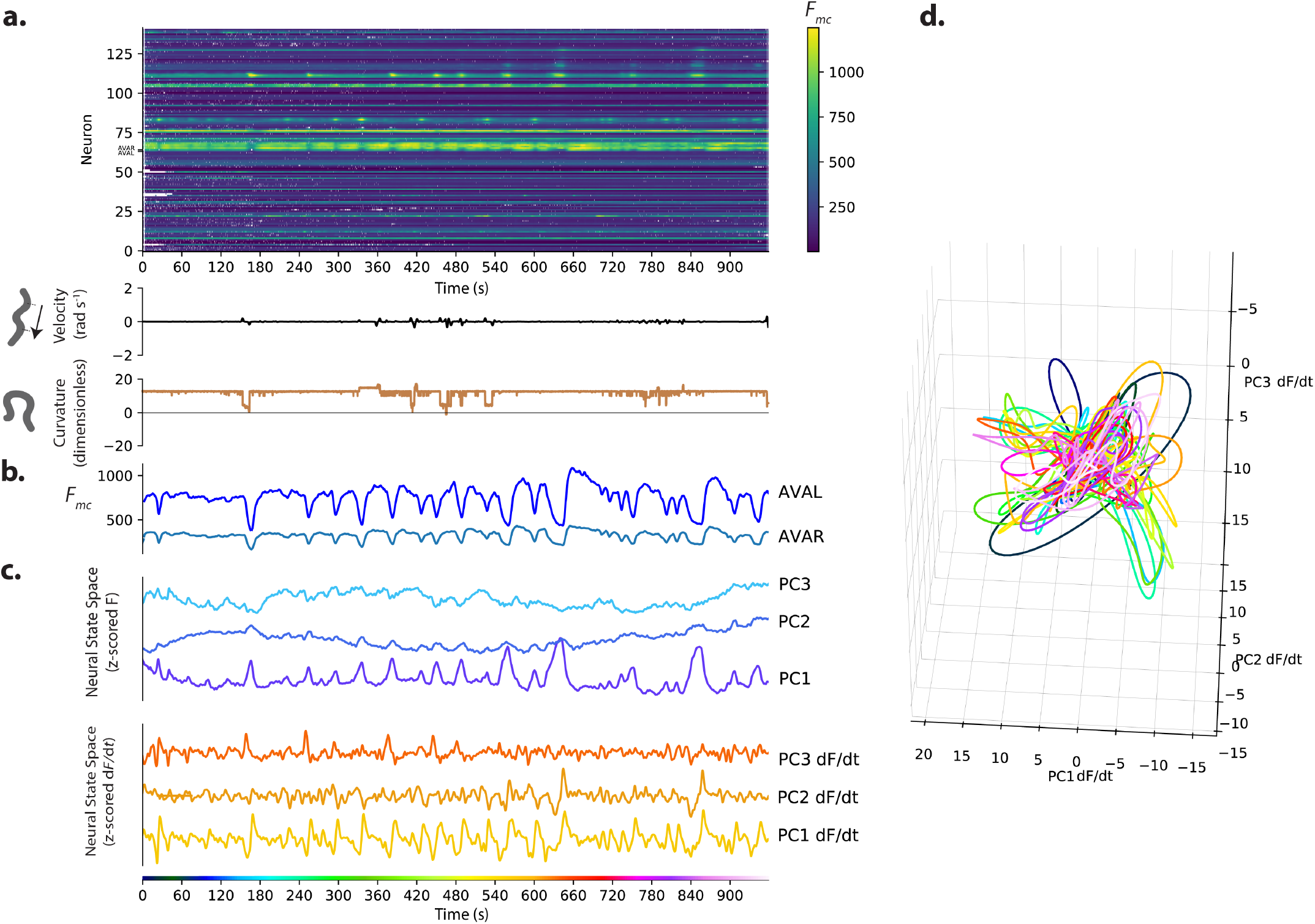
Calcium activity is recorded from an animal immobilized with nano-beads, recording AML310_G . a.) Calcium activity. b.) Activity of neurons AVAL and AVAR. c.) Population activity (or its temporal derivative) from (a) is shown projected onto its first three PCs, as determined by only the immobilized portion of the recording. d.) Neural state space trajectories from (b) are plotted in 3D.

## Notes

https://doi.org/10.17605/OSF.IO/R5TB3

### Competing Interest Statement

The authors have declared no competing interest.

### Summary of Updates

Major revision with additional new experiments. In particular, we include extensive new measurements of neuron pair AVA that demonstrate that our population recordings are consistent with prior measurements of single neurons.

